# CD4^+^ T cells promote fibrosis during metabolic dysfunction-associated steatohepatitis

**DOI:** 10.1101/2025.06.27.660220

**Authors:** Lucía Valenzuela-Pérez, Hyun Se Kim Lee, Rachel L. Bayer, Shravan K. Mishra, Alexander M. Washington, Qianqian Guo, Adam Herman, Rondell P. Graham, Malaz M. Sidahmed, Edward Ssali, Adna A. Hassan, Ece Janet Dinc, Kevin D. Pavelko, Gregory J. Gores, Patrick Starlinger, Xavier S. Revelo, Samar H. Ibrahim, Enis Kostallari, Adebowale O. Bamidele, Petra Hirsova

**Affiliations:** Division of Gastroenterology and Hepatology, Mayo Clinic, Rochester, Minnesota, USA; Mayo Clinic Graduate School of Biomedical Sciences, Mayo Clinic, Rochester, Minnesota, USA; Department of Surgery, University of Minnesota, Minneapolis, Minnesota, USA; Department of Laboratory Medicine and Pathology, Mayo Clinic, Rochester, Minnesota, USA; Immune Monitoring Core, Mayo Clinic, Rochester, Minnesota, USA; Division of Hepatobiliary and Pancreas Surgery, Mayo Clinic, Rochester, Minnesota, USA; Centre of Physiology and Pharmacology, Medical University of Vienna, Vienna, Austria; Department of Integrative Biology and Physiology, University of Minnesota, Minneapolis, Minnesota, USA; Center for Immunology, University of Minnesota, Minneapolis, Minnesota, USA; Division of Pediatric Gastroenterology and Hepatology, Mayo Clinic, Rochester, Minnesota, USA; Department of Biochemistry and Molecular Biology, Mayo Clinic, Rochester, Minnesota, USA; Department of Immunology, Mayo Clinic, Rochester, Minnesota, USA

**Keywords:** NAFLD, NASH, MASLD, MASH, T cells, subsets, western diet

## Abstract

Unresolved inflammation and fibrosis are the two key features of metabolic dysfunction-associated steatohepatitis (MASH), a progressive form of steatotic liver disease that can evolve into cirrhosis and liver cancer. Although innate immunity has been well studied in MASH, the role of CD4⁺ T cells remains underexplored despite their potential to coordinate immune responses by providing help to other immune cells, promoting inflammation, or regulating immune activity through effector and regulatory subsets. To better understand the role of CD4^+^ T cells in the pathogenesis of MASH, we comprehensively characterized hepatic CD4^+^ T cells in murine and human MASH at a single-cell protein, transcriptional, and functional level. Mass cytometry and CITE-sequencing revealed a marked shift in intrahepatic CD4⁺ T-cell composition in MASH, with enrichment of Th1, regulatory, and cytotoxic CD4⁺ T cells. Similar phenotypic changes were mirrored in the peripheral blood and validated in human MASH samples. Functional assays demonstrated increased production of IFNγ and TNFα by hepatic CD4⁺ T cells, highlighting their proinflammatory effector activity. Transcriptomic profiling identified Tnfrsf4 (OX40) upregulation in hepatic CD4⁺ T cells during MASH. Therapeutic blockade of the OX40L-OX40 axis reversed hepatic fibrosis and improved histologic disease scores in mice with established MASH, and also decreased inflammatory markers in a human *ex vivo* liver model. Together, these studies provide a proteogenomic single-cell atlas for hepatic CD4⁺ T cells and uncover a CD4⁺ T cell-dependent immunopathogenic circuit as a promising immunotherapeutic target to alleviate MASH and liver fibrosis.

## INTRODUCTION

Metabolic dysfunction-associated steatotic liver disease (MASLD), formerly known as nonalcoholic fatty liver disease, is an umbrella term encompassing a range of liver conditions characterized by excessive hepatic lipid accumulation. As the most prevalent chronic liver disease globally, MASLD is tightly linked to metabolic disorders and obesity-associated comorbidities such as type 2 diabetes.^1^ While initially asymptomatic, MASLD can progress to the inflammatory and fibrotic form termed metabolic dysfunction-associated steatohepatitis (MASH). MASH, a progressive condition, is marked by hepatocyte ballooning and cell death, and tissue inflammation in both the parenchymal and portal areas, leading to varying degrees of fibrosis.^2^ The progression from isolated steatosis to MASH is driven by complex interactions between lipid toxicity, oxidative stress, and inflammation, with the development of liver fibrosis being the main predictor of morbidity and mortality in patients with MASH.^3^ If left untreated, MASH can progress to cirrhosis and significantly increase the risk of developing hepatocellular carcinoma, which is the most rapidly increasing indication for liver transplantation in the United States.^4^ While current therapeutic options for MASH are limited, the recent approval of Resmetirom provides the first pharmacotherapy for MASH, albeit a significant proportion of MASH patients do not respond.^5^ This underscores the critical need to further investigate cellular and molecular mechanisms driving disease progression, particularly in the context of liver inflammation, which can complement targeting metabolic aspects of MASH.

Inflammation plays a pivotal role in the transition from isolated steatosis to MASH, and emerging evidence highlights the critical involvement of the adaptive immune system in this process. Although innate immune responses are traditionally considered the primary initiators of MASH, the adaptive immune response and its ability to amplify the inflammatory milieu have gained increasing attention as a key mechanism in the pathogenesis of the disease and its progression to fibrosis and liver cancer. Infiltration of conventional T lymphocytes into the liver parenchyma is increasingly recognized as a hallmark of MASH, and the modulation of T-cell recruitment or function has been shown to impact MASH severity in preclinical models.^6–8^ However, a deep understanding of CD4^+^ T-cell phenotype and function in MASH has been limited, despite their potential to orchestrate immune responses in inflamed tissues. Egressing from the thymus to peripheral lymphoid sites, CD4^+^ T cells are crucial for immune responses, orchestrating tissue homeostasis and amplifying pathology in a context-dependent manner. Upon activation, naïve CD4^+^ T cells differentiate into various subsets defined by specific cytokine profiles and transcription factor expression, including Th1 (IFNγ, T-bet), Th2 (IL-4, GATA3), Th17 (IL-17, RORγT), and regulatory T (Treg; IL-10/TGFβ, FoxP3) cells.^9^ However, the precise hepatic CD4^+^ T-cell landscape in MASH and the mechanisms by which CD4+ T cells may promote disease progression remain poorly defined, hindering immuno-therapeutic discoveries for MASH treatment.

Herein, we provide detailed insights into heterogeneous hepatic CD4^+^ T-cell populations in murine and human MASH using a combination of single-cell analyses as well as functional evidence for the role of CD4^+^ T cells in murine and human models. The absence of CD4^+^ T cells in murine MASH significantly reduces hepatic injury, inflammation, and fibrosis, independent of insulin resistance or liver steatosis. Through single-cell approaches, we identify significant changes in the heterogeneity of hepatic CD4^+^ T-cell subsets during MASH, characterized mainly by the expansion of Th1, Treg, and cytotoxic populations in both the liver and peripheral blood. Thus, this study provides a useful resource for a common and human disease-relevant MASH model, often used to study disease pathogenesis and test therapeutic approaches. We also identified the OX40 ligand (OX40L)-OX40 axis as a key immunoregulatory pathway in MASH, demonstrating that inhibition of the OX40L-OX40 signaling reverses hepatic fibrosis in established murine MASH. Additionally, treatment of human precision-cut liver slices (PCLS) with Rocatinlimab, a human OX40 antagonistic antibody, reduced markers of inflammation. Together, these findings provide new insights into the role of CD4^+^ T cells in MASH and validate the potential for CD4^+^ T cell-directed immunotherapies.

## MATERIALS AND METHODS

## RESULTS

### CD4 deletion attenuates steatohepatitis and liver fibrosis in a murine MASH model

To broadly examine the role of CD4^+^ T cells in MASH development, we utilized CD4 whole-body knockout (CD4^-/-^) mice, which lack CD4^+^ T cells due to the absence of the CD4 co-receptor, resulting in a developmental block during thymic positive selection. CD4^-/-^ mice and wild-type (WT) controls were fed a diet high in fat, fructose, and cholesterol (FFC) for 24 weeks (Fig. 1A). The FFC diet is a well-validated and human disease-relevant MASH model with cardiometabolic comorbidities of obesity, insulin resistance, adipose tissue inflammation, and steatohepatitis with fibrosis.^10,11^ Given that metabolic status can influence MASH development, we assessed key metabolic parameters in our mice. After 20 weeks of feeding, mice were subjected to comprehensive laboratory animal monitoring, including measurements of body composition and metabolic parameters. CD4 deletion did not affect the fat and lean mass of mice on the FFC diet (Fig. S1A). Food intake, body weight, and metabolic parameters, such as metabolic rate and energy expenditure, were not different between FFC-fed CD4^-/-^ and WT mice (Fig. S1B-F). Insulin resistance, assessed after 22 weeks of feeding by HOMA-IR, was significantly increased in FFC-fed WT mice compared to chow controls, but no difference was observed between the FFC-fed CD4^-/-^ and WT mice (Fig. 1B, Fig. S1G). FFC-fed CD4^-/-^ mice developed the same level of obesity as FFC-fed WT mice (Fig. 1C, Fig. S1H). Interestingly, although the body weight and adipose weight were not affected by CD4 deletion (Fig. S1I), the liver-to-body weight ratio decreased in FFC-fed CD4^-/-^ mice (Fig. 1D), likely secondary to reduced immune cell infiltration. Plasma alanine aminotransferase (ALT) and aspartate aminotransferase (AST) activity, markers of hepatocyte injury, significantly decreased in FFC-fed CD4^-/-^ compared to FFC-fed controls (Fig. 1E-F). In further phenotypic studies, hepatic steatosis was not different between FFC-fed CD4^-/-^ and WT mice, as evident from H&E staining and biochemical quantification of liver triglycerides (Fig. 1G-H). Since hepatic accumulation of monocyte-derived macrophages is a hallmark of MASH, we evaluated their presence *in situ* using immunohistochemistry. FFC feeding increased the abundance of Lgals3^+^ (a marker of phagocytically active macrophages) and F4/80^+^ (a general macrophage marker) cells in the liver, which was significantly reduced in FFC-fed CD4^-/-^ mice (Fig. S1J). Given that chronic liver injury and inflammation in MASH drive fibrogenesis, we next assessed the extent of liver fibrosis in our cohort. Sirius red staining for collagen deposition and immunopositivity for alpha-smooth muscle actin (αSMA), a marker of activated hepatic stellate cells, were both elevated in FFC-fed WT mice and markedly reduced in CD4^-/-^ counterparts (Fig. 1I-L). Histological scoring by a blinded liver pathologist confirmed significantly attenuated fibrosis in FFC-fed CD4^-/-^ mice compared to WT controls (Fig. 1M). Fibrosis in MASH is linked to a profibrogenic macrophage population expressing the "Fab5" gene signature: TREM2, CD9, FABP5, osteopontin (SPP1), and GPNMB.^12^ To investigate whether CD4⁺ T cells influence this macrophage phenotype, we performed gene expression analysis on flow-sorted hepatic monocyte-derived macrophages (CD45⁺ Ly6G⁻ CD11b⁺ F4/80⁺) from FFC-fed CD4^-/-^ mice and WT controls (Fig. S1K). All Fab5 genes were significantly downregulated in macrophages from CD4^-/-^ livers (Fig. 1N), suggesting that CD4^+^ T cells promote the emergence of profibrogenic macrophages in MASH. Together, these results demonstrate that the absence of CD4⁺ T cells reduces liver injury, inflammation, and fibrosis in the context of FFC-induced MASH, without impacting systemic metabolic dysfunction.

**Figure 1:**
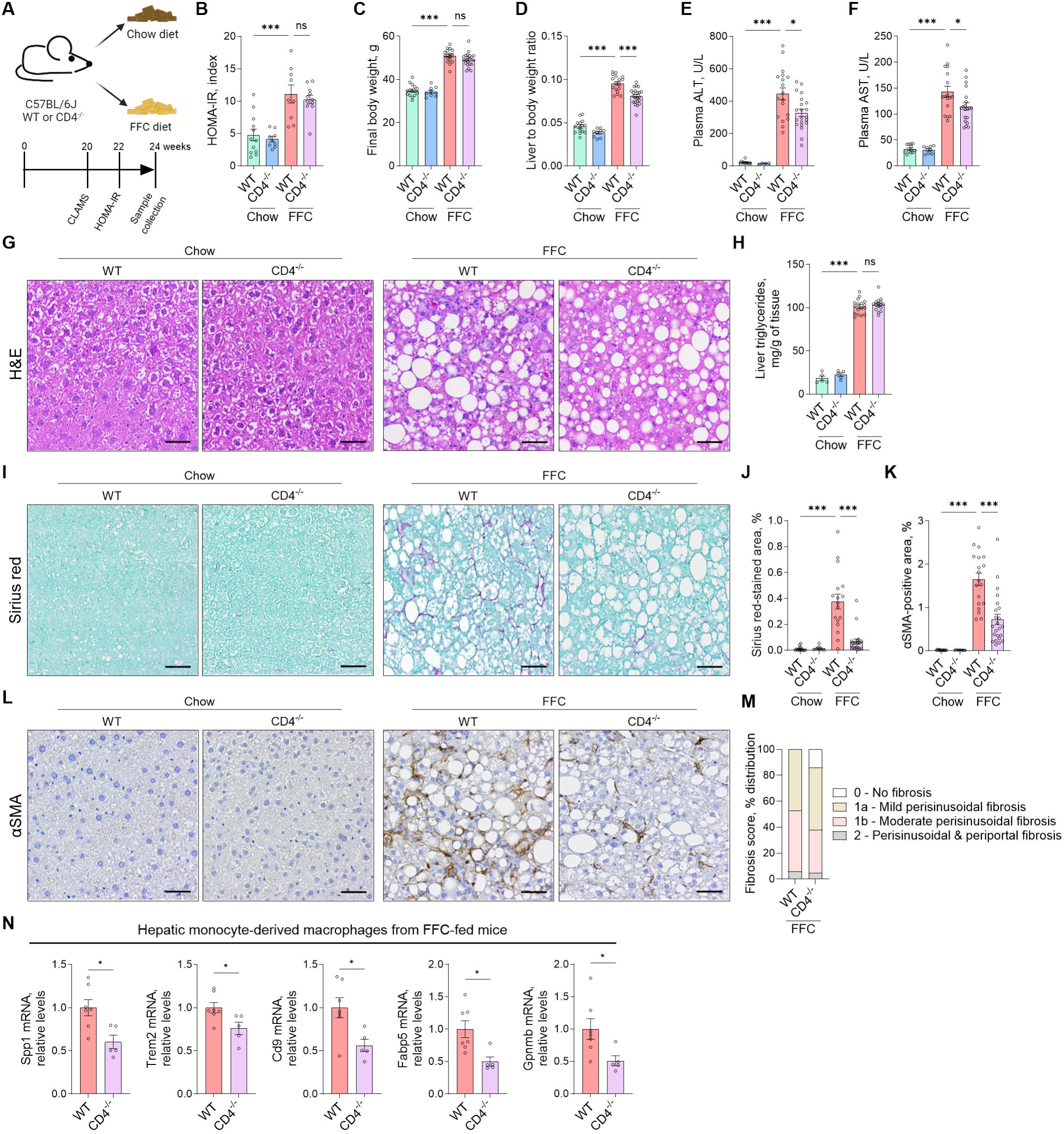
Liver injury and fibrosis are decreased in FFC diet-fed CD4^-/-^ mice. **(A)** C57Bl/6J (WT) and CD4 knockout (CD4^-/-^) mice were fed a chow or FFC diet for 24 weeks. **(B)** Insulin resistance was measured by HOMA-IR using fasting glucose and fasting insulin values at week 22. **(C)** Body weight at the time of sacrifice. **(D)** Liver weight normalized to the body weight. **(E-F)** Liver injury assessed by plasma ALT and AST activity. **(G)** H&E staining on liver tissue sections. **(H)** Liver triglycerides quantification. **(I-J)** Sirius red staining for liver collagen deposition and quantification **(K-L)** Immunohistochemistry and quantification for liver αSMA, a marker of activated hepatic stellate cells. Representative images taken with a 20X objective are shown; the scale bar is 50 µm. **(M)** Fibrosis scoring by a blinded pathologist. **(M)** Gene expression by qPCR in flow-sorted hepatic monocyte-derived macrophages from FFC-fed mice. N of samples indicated in the graphs. *p < 0.05, ***p < 0.001. ns, non- significant. Abbreviations: WT, wild-type; FFC, high fat, fructose, and cholesterol; HOMA-IR, Homeostatic Model Assessment for Insulin Resistance; ALT, alanine aminotransaminase; AST, aspartate aminotransferase; H&E, Hematoxylin and eosin; αSMA, alpha-smooth muscle actin.

### Immunophenotyping of hepatic CD4^+^ T cells reveals significant enrichment in Th1, cytotoxic, and Treg cells in murine MASH

The frequency of hepatic CD4^+^ T cells in all liver leukocytes (CD45^+^ cells) is not markedly different between chow and FFC-fed mice (Fig. S2A), suggesting that the phenotype of these cells, rather than numbers, is more important for disease development. CD4^+^ T cells differentiate into multiple sublineages or subsets with distinct functions. Despite this heterogeneity, hepatic CD4^+^ T-cell phenotypes have not yet been comprehensively characterized in murine MASH. To this end, we analyzed the hepatic CD4^+^ T-cell composition in FFC-fed mice by mass cytometry by time-of-flight (CyTOF), which enabled us to simultaneously assess a total of 44 well-established protein markers at a single-cell level. First, mice were fed a chow or MASH-inducing FFC diet for 24 weeks, followed by isolation of intrahepatic CD4^+^ T cells (Fig. 2A). CD4^+^ T cells were then labeled with rare-earth metal antibodies and immunophenotyped by mass cytometry. We gated on CD3^+^ TCRβ^+^ CD4^+^ CD8^-^ cells to perform the downstream analysis on the pure CD4^+^ T-cell population. We identified 16 clusters of CD4^+^ T cells, including naïve/central memory cells, various T helper (Th) subsets such as Th1, Th17, and T follicular helper cells (Tfh), regulatory T cells (Treg), and cytotoxic CD4^+^ cells, as shown in the tSNE plot (Fig. 2B). One additional cluster represented cell doublets (cluster 12, ∼0.5% of analyzed cells). Each diet group exhibited a distinct composition of CD4^+^ T-cell population, as evidenced in the tSNE plots of cell densities and heatmap displaying the relative cluster frequencies and unsupervised hierarchical clustering of each sample (Fig. 2C-D). The 16 clusters of CD4 T cells were annotated based on the relative expression of proteins that are well-established markers for specific sublineages (Fig. 2E-F, Fig. S2B-E). In the FFC diet, the most abundant clusters were clusters 17, 10, and 14, all characterized by high expression of CD44 and T-bet, a lineage-defining transcription factor of Th1 cells (Fig. 2G). In addition, these clusters had different expression levels of IL7R and PD1, suggesting these clusters represent different stages of activation - from early (IL7R^+^ PD1^-^) to late (IL7R^-^ PD1^+^) activation (Fig. 2G). Cluster 16 was another abundant cluster in the MASH-inducing diet, which was also high in T-bet, but the expression of Eomes and PD1 indicated that this cluster has a cytotoxic profile (Fig. 2H). FFC diet also promoted accrual of cells expressing FoxP3, (a transcription factor essential for the development, function, and stability of Tregs) such as FoxP3^+^ CD25^high^ Tregs and FoxP3^+^ CD25^low^ cells (Fig. 2I). We detected two clusters of Th17 cells, which differ from each other by the expression of CD103 (integrin αE) and integrin β7. The frequency of hepatic Th17 cells did not change by FFC feeding (Fig. 2J). In contrast, in chow mice, the most prevalent clusters were naïve/central memory CD4^+^ T cells (clusters 4, 5, 6, and 11), defined as CD62L^+^ CCR7^+^ CD44^-/+^ cells (Fig. S2C, E). These data demonstrate a highly distinct CD4^+^ T-cell composition between healthy and MASH livers. A unique effector CD4^+^ T-cell pattern with Th1, cytotoxic, and Treg cell enrichment in MASH livers provides novel implications for disease pathogenesis.

**Figure 2.**
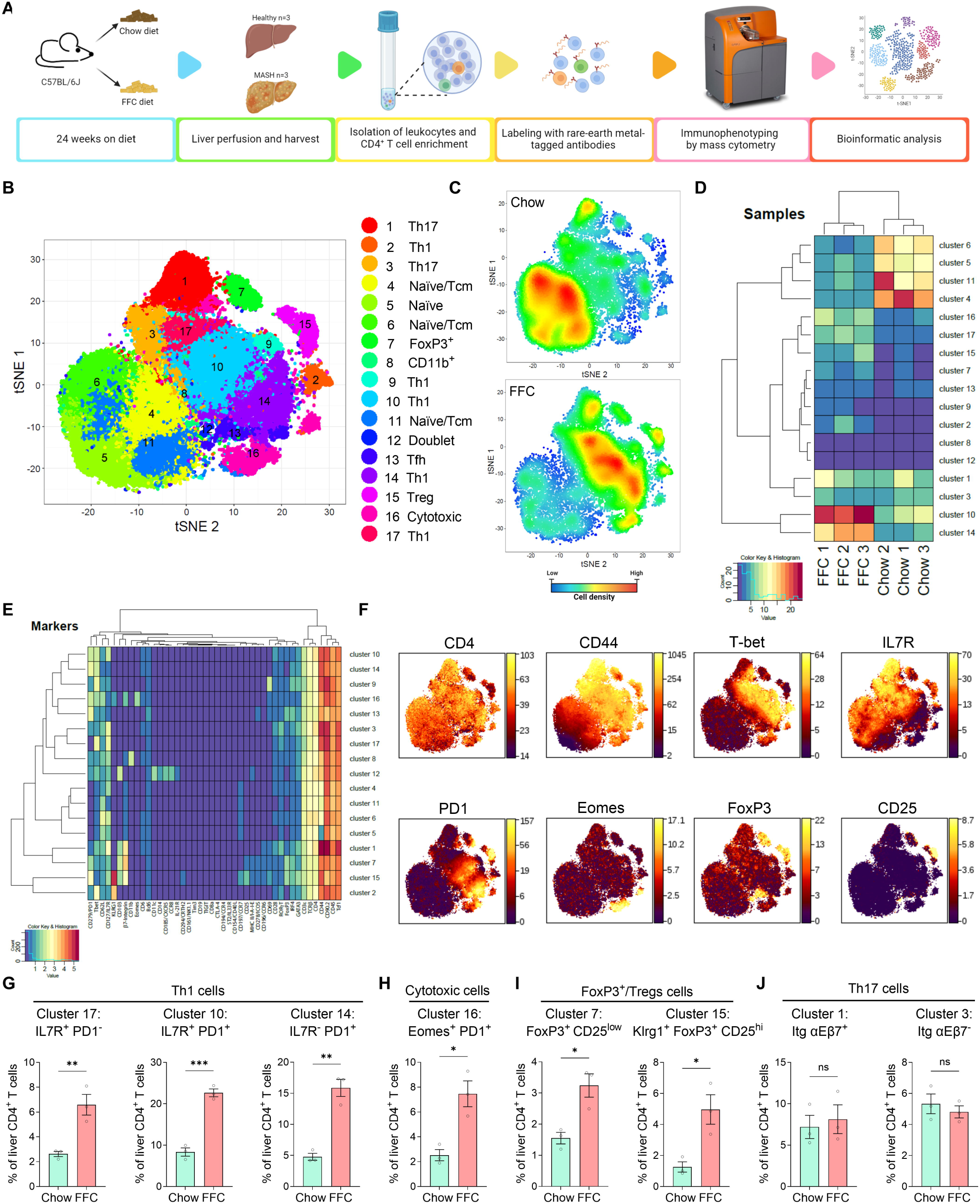
Murine MASH livers show significant enrichment in Th1, cytotoxic, and Foxp3^+^ cells. **(A)** Workflow for immunophenotyping of hepatic CD4^+^ T cells by mass cytometry (CyTOF). **(B)** Seventeen unique clusters of cells were identified using an R-phenograph clustering algorithm, visualized on a tSNE plot, and annotated. **(C)** A tSNE plot of relative cell densities. **(D)** Unsupervised hierarchical clustering of chow and FFC samples based on the relative abundance of CD4^+^ T-cell clusters. **(E)** A heatmap of the relative intensity of 44 protein markers across identified clusters with unsupervised hierarchical clustering. **(F)** Representative heat maps of markers relevant for clusters enriched in FFC samples. **(G-J)** Frequencies of Th1, cytotoxic CD4^+^, Foxp3^+^, and Th17 cell clusters. N of samples indicated in the graphs. *p < 0.05, **p < 0.01, ***p < 0.001, ns, non-significant. Abbreviations: FFC, high fat, fructose, and cholesterol; CyTOF, cytometry by time-of-flight; tSNE, t-distributed stochastic neighbor embedding.

### Peripheral blood and liver share similar changes in the CD4^+^ T-cell landscape during MASH

Finding that hepatic CD4^+^ T cells have distinctive phenotypes in MASH compared to normal liver, we next examined whether this is a unique feature of the liver tissue or if it is also reflected in the peripheral blood, as identifying shared signatures could enable noninvasive monitoring of liver immune dysregulation. To this end, peripheral blood mononuclear cells (PBMCs) from chow and FFC-fed mice were subjected to mass cytometry using an antibody panel for T-cell markers (Fig. 3A). After gating on CD4^+^ T cells (defined as CD45^+^ CD3^+^ TCRβ^+^ CD4^+^ CD8^-^ cells), we detected 18 distinct CD4^+^ T-cell clusters, which included naïve/Tcm cells, Th1, cytotoxic, Treg, and Th17 cells (Fig. 3B-C, Fig. S3). Similar to the liver, circulating CD4^+^ T cells in FFC-fed mice exhibited a marked enrichment in select subsets, as evidenced by tSNE plots of cell densities and a heatmap of relative cluster frequencies (Fig. 3D-E). In particular, clusters increased in MASH mice compared to chow were represented by Th1, cytotoxic, Treg, and Th17 cells (Fig. 3F). Comparing the CD4^+^ T-cell changes during MASH, liver and peripheral blood show similar fold increases in Th1, Treg, and cytotoxic CD4^+^ T cells. Thus, phenotypic shifts in the CD4^+^ T-cell population during MASH are detectable in both the liver and circulation and may potentially serve as a noninvasive blood-based biomarker for MASH.

**Figure 3.**
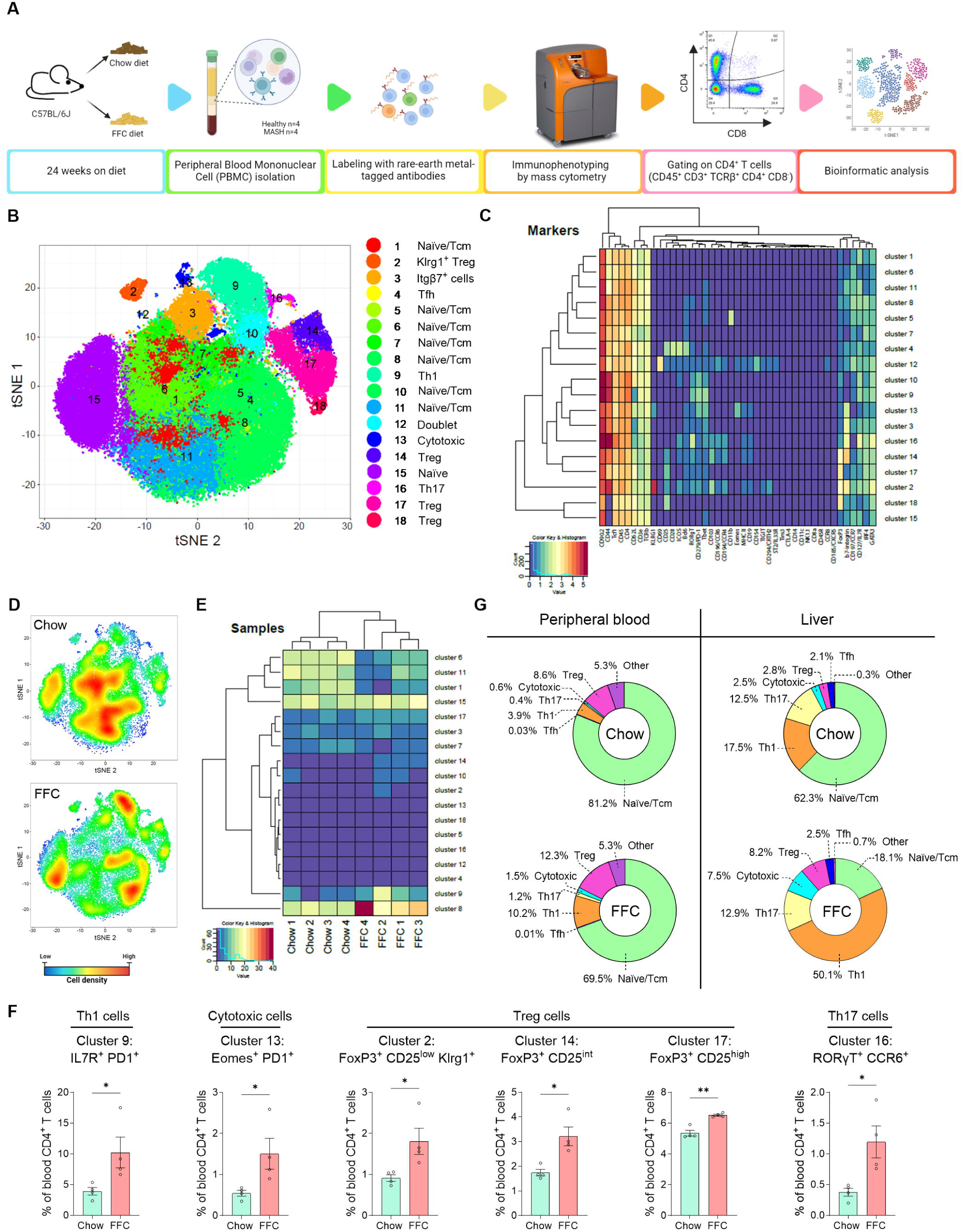
Peripheral blood and liver share similar changes in the CD4^+^ T cell landscape during MASH. **(A)** Workflow for immunophenotyping CD4^+^ T cells in peripheral blood by mass cytometry (CyTOF). **(B)** CD4^+^ T cell clusters by tSNE plot R-phenograph. **(C)** A heatmap of the relative intensity of protein markers across identified clusters with unsupervised hierarchical clustering. **(D)** A tSNE plot of relative cell densities. **(E)** Unsupervised hierarchical clustering of chow and FFC samples based on the relative abundance of CD4^+^ T cell clusters. **(F)** Frequencies of Th1, cytotoxic, Treg and Th17 cells. **(G)** Comparison of CD4^+^ T cell subset between peripheral blood and liver. Relative abundance of Th1, cytotoxic CD4^+^, and Treg cell clusters. N of samples indicated in the graphs. *p < 0.05, **p < 0.01, ***p < 0.001, ns, non-significant. Abbreviations: FFC, high fat, fructose, and cholesterol; PBMC, peripheral blood mononuclear cell; CyTOF, cytometry time-of-flight; tSNE, t-distributed stochastic neighbor embedding.

### Single-cell atlas of intrahepatic CD4^+^ T cells in healthy and MASH livers

Since CyTOF-identified subsets are influenced by the antibody selection, we next aimed to characterize the hepatic CD4^+^ T-cell landscape in an unbiased manner based on RNA transcriptome. To this end, we performed cellular indexing of transcriptomes and epitopes by sequencing (CITE-seq), which allows for RNA and proteins to be analyzed simultaneously at a single-cell level. FACS-sorted CD4^+^ T cells from the liver of chow-fed and FFC-fed mice were labeled with a panel of oligo-conjugated antibodies (antibody-derived tags/ADT) against 17 relevant T-cell surface markers and subjected to droplet-based 3’-end 10x Genomics scRNA-seq (Fig. 4A). Unsupervised clustering and Uniform Manifold Approximation and Projection (UMAP) identified and visualized 15 distinct clusters of hepatic CD4^+^ T cells based on their distinct transcriptional profile (Fig. 4B-C, Fig. S4A-B). A combination of the protein and marker gene expression was then used for cluster annotation (Fig. 4D-E, Fig. S4C-D). A clear difference was evident in the relative distribution of chow and FFC-derived hepatic CD4^+^ T cells among the 15 clusters, indicating again strong phenotypic changes to the CD4^+^ T-cell population during MASH. We identified four subsets representing naïve/central memory (Tcm) CD4^+^ T cells expressing *Sell* (CD62L protein)*, Ccr7* (CCR7 protein), and *Lef1*, which were enriched in chow livers (cluster 2, 3, 5, and 9; Fig. 4F-G). The most abundant CD4^+^ T-cell subset in FFC liver (cluster 0) corresponded to cells with Th1 polarization, expressing markers such as *Tbx21*, *Ifng*, *Ifngr1*, *Stat4*, *Cxcr3* and activation markers such as *Ly6a*, *Icos*, *Tnfsf18*, *Il18r1*, *Cd44* (CD44 protein), *Tnfrsf4*, and *Slamf6*. The second largest cluster (cluster 1), also enriched in FFC, represented cytotoxic/tissue resident CD4^+^ T cells, displaying cytotoxic gene signatures (*Nkg7, Eomes, Gzmk, Gzmb, Prf1, Fasl, Ccl5, Ccl4, Cd160*) and the tissue residency markers *Cxcr6* and CXCR6 (protein). Other clusters increased in FFC compared to chow livers (clusters 4, 8, and 10) were Tregs (*Foxp3*, *Ikzf2*, *Il2ra* [CD25 protein], NRP1 and CD103 protein), Th17 (*Rorc, Il17a, Il23r,* CCR6 protein) and cells with IFN response genes (Isg15, Ifit1). We also identified two clusters of cycling cells and a cluster of Th2 (*Gata3, Pparg, Il1rl1*) and Tfh (*Bcl6, Il21, Cxcr5/*CXCR5 protein) cells, which represented minor populations of intrahepatic CD4^+^ T cells. Together, the novel subsets described herein are represented mainly by cytotoxic CD4^+^ T cells, IFN response cells, FoxP3^+^ CD25^neg^ cells, and Tfh cells, which provide help to B cells in secondary lymphoid organs and are not expected to be present in liver tissue. Next, we assessed the developmental fate of CD4^+^ T cells by applying trajectory analysis to the transcriptomic data. Defining the naïve CD4^+^ T cells as the origin, several major inferred cellular trajectories towards the main differentiated subsets were observed (Fig. 4H). For example, the data indicate a common developmental trajectory for Tregs and Th17 cells or Th1 and cytotoxic CD4^+^ T cells.

Finally, to assess the conservation of CD4⁺ T cell states between species, we compared the CD4^+^ T cell landscape between murine and human MASH. To this end, we used label transfer and projection techniques and mapped the human CD45RA^-^ CD4^+^ T cell transcriptome from MASH patients (GEO: GSE217235) onto our murine scRNAseq reference dataset. We detected the presence of similar clusters in human MASH hepatic CD4^+^ T cells, with a high proportion of Th1 cells, especially (Fig. 4I-J). Since the scRNAseq was done on memory CD4^+^ T cells, the corresponding clusters of naïve cells were largely missing. To further investigate the CD4^+^ T cell subsets in a spatial context, we performed *in situ analysis* of Th1, Tregs, and cytotoxic CD4⁺ T cells in human liver tissue using multiplex spatial phenotyping with the PhenoCycler-Fusion system. Liver sections from patients with isolated steatosis or MASH were stained with a panel of antibodies against select T-cell markers, including T-bet, Eomes, FoxP3, and RORγt. Given the potential presence of portal inflammation in MASH (Fig. S4E), we focused our analysis on CD4^+^ T cells (defined as CD45^+^ CD3^+^ CD4^+^ CD8^-^) localized within the lobular parenchyma. All four CD4⁺ T cell subsets, T-bet⁺ (Th1), Eomes⁺ (cytotoxic), FoxP3⁺ (Tregs), and RORγt⁺ (Th17), were elevated in MASH livers compared to isolated steatosis, with T-bet⁺ CD4⁺ T cells showing a statistically significant increase (Fig. 4K). This spatial immunophenotyping aligns with our transcriptomic and mass cytometry data, reinforcing the enrichment of Th1, Treg, and cytotoxic CD4^+^ T cells in the MASH livers. Altogether, the transcriptome-based unbiased results and spatial phenotyping corroborate our mass cytometry data and demonstrate the enrichment of Th1, Tregs, and cytotoxic CD4^+^ T cells in MASH livers. Similarities between murine and human MASH CD4^+^ T cells underscored the relevance of Th1 cells in the pathogenesis of the disease.

**Figure 4.**
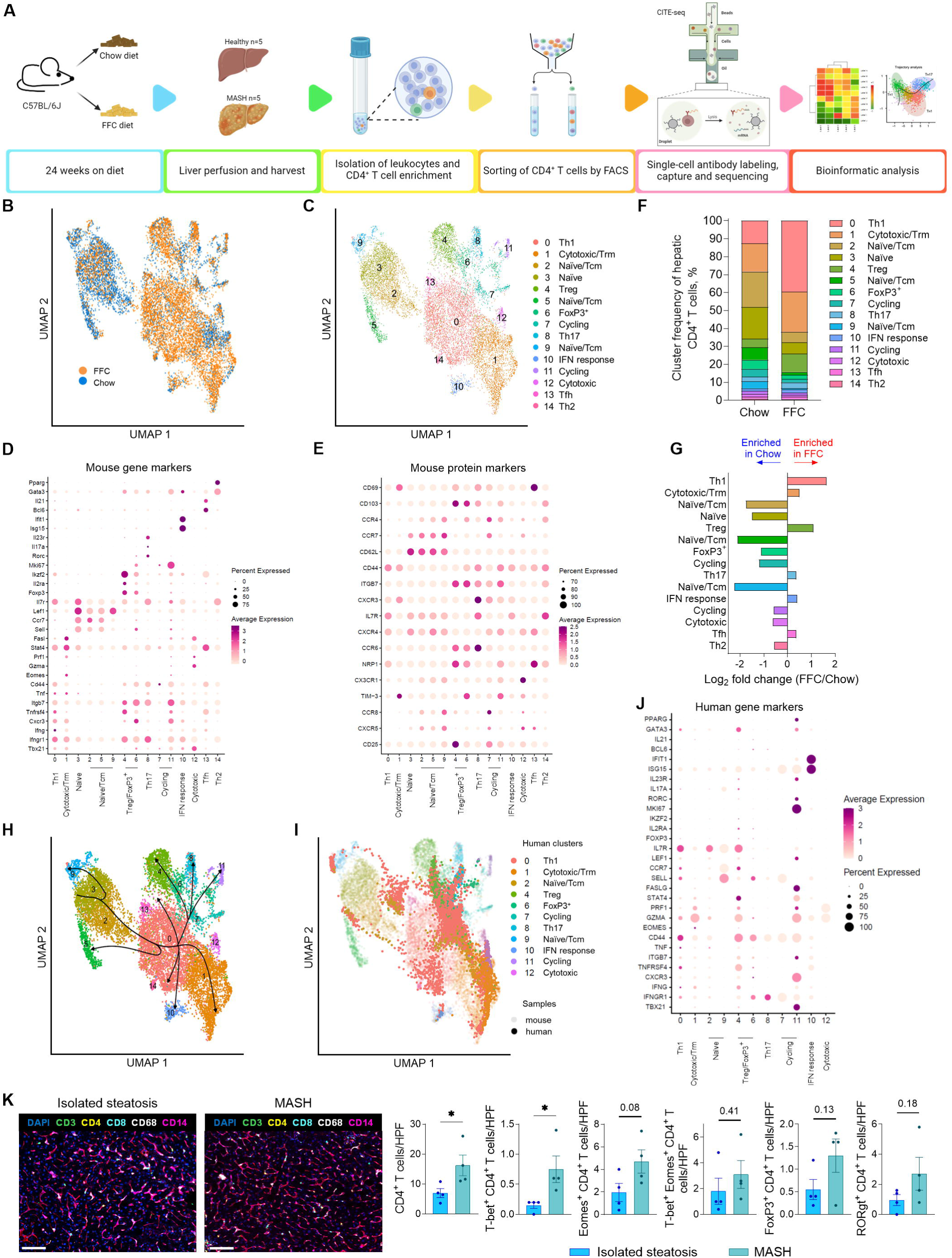
Single-cell atlas of intrahepatic CD4 T cells in healthy and MASH liver. **(A)** Workflow for CITEseq in hepatic CD4^+^ T cells. **(B)** UMAP projection of 12,701 hepatic CD4^+^ T cells from chow and FFC-fed mice. **(C)** UMAP projection of 15 distinct hepatic CD4^+^ T-cell clusters. **(D)** Average gene expression of select markers. **(E)** Relative protein marker expression based on sequencing of antibody-derived tags. **(F)** Cluster frequencies of hepatic CD4^+^ T cells. **(G)** Log fold change of cluster abundance. **(H)** Trajectory analysis. **(I)** UMAP of human hepatic CD4^+^ T cells from MASH patients (n=3) projected on murine intrahepatic CD4^+^ T-cell clusters. **(J)** Average gene expression for select human markers. **(K)** Representative images of human liver samples using Phenocycler-Fusion phenotyping and quantification of CD4^+^ T-cell subsets per high-power field. N of samples indicated in the graphs. *p < 0.05 or p value as shown. Abbreviations: FACS, fluorescence-activated cell sorting; CITE-seq, Cellular Indexing of Transcriptomes and Epitopes sequencing; UMAP, Uniform Manifold Approximation and Projection; HPF, high power field.

### Hepatic CD4 T cells in MASH increase proinflammatory cytokine secretion

In both CyTOF and CITEseq data, we detected increased numbers of effector CD4^+^ T cells (CD62L^-^ CCR7^-^ CD44^hi^) in MASH livers compared to healthy livers. In addition, gene set enrichment analysis of scRNAseq data indicated that cytokine production is increased in various subsets of CD4^+^ T cells (Fig. S5A). To confirm the effector function, we assessed the gene and protein expression of select effector cytokines. To overcome the lower sequencing depth of scRNAseq, we measured mRNA expression with a targeted approach using NanoString nCounter technology, providing a copy number of transcripts following hybridization with target-specific probes. Among the top 5 upregulated transcripts in freshly isolated hepatic CD4^+^ T cells in MASH livers were those encoding IFNγ and TIM3, a surface marker of IFNγ-producing cells (Fig. S5B). To assess cytokine production at the protein level, isolated hepatic CD4^+^ T cells were cultured in the presence of brefeldin A (to block protein secretion) with or without concurrent stimulation by phorbol 12-myristate 13-acetate (PMA) and ionomycin (Fig. 5A). Flow cytometric analysis revealed that FFC-derived hepatic CD4^+^ T cells produced more IFNγ, which was further accentuated by PMA/ionomycin stimulation (Fig. 5A, Fig. S5C). Stimulated IFNγ^+^ CD4^+^ T cells also produced higher amounts of TNFα. IL-17A^+^ CD4^+^ T cells were only slightly increased by the stimulation. IFNγ, TNFα, IL17A, and other cytokines were also detected in the cell culture media when hepatic CD4^+^ T cells from chow and FFC mice were re-stimulated *ex vivo* in the absence of brefeldin A (Fig. S4D). Altogether, these data demonstrate an increased IFNγ and TNFα production in hepatic CD4^+^ T cells in MASH and are consistent with the enrichment of Th1 cells, potentially promoting disease pathogenesis.

**Figure 5.**
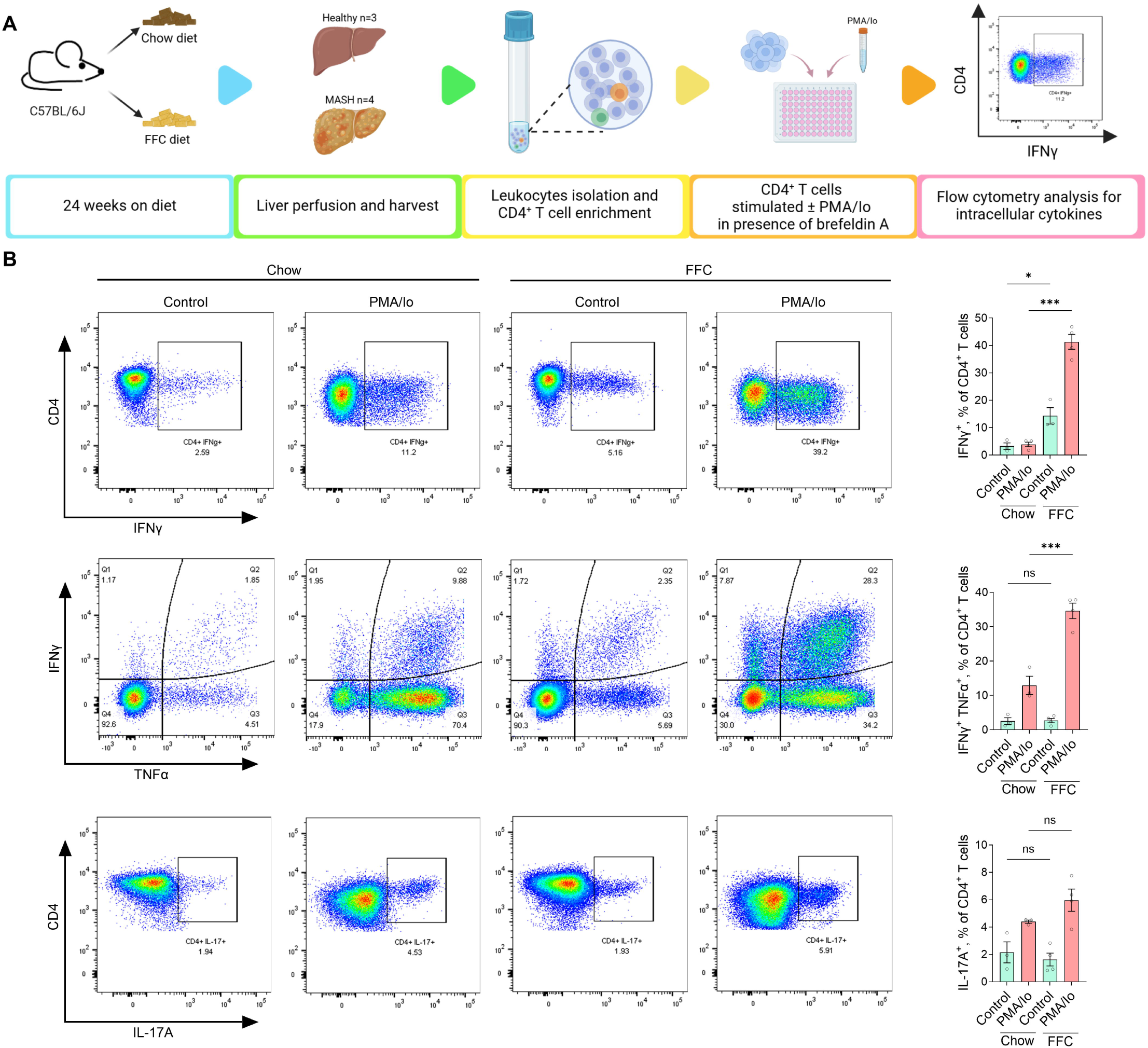
Liver CD4^+^ T cells in MASH display increased production of proinflammatory cytokines. **(A)** Workflow for analysis of intracellular cytokines. **(B)** CD4^+^ T cells were cultured *ex vivo* with or without stimulation with PMA+ionomycin for 4 hours in the presence of brefeldin A. Intracellular cytokines were quantified by flow cytometry. N of samples indicated in the graphs. *p < 0.05, ***p < 0.001. ns, non- significant. Abbreviations: PMA/Io, phorbol 12-myristate 13-acetate/ionomycin.

### Tnfrsf4/OX40 is upregulated in CD4^+^ T cells in MASH

Considering that MASH livers have distinct CD4^+^ T-cell phenotypes, we next searched for a potential MASH-associated molecular target that could be therapeutically manipulated. We analyzed our scRNAseq dataset for differential gene expression between FFC and chow cells at the pseudo-bulk level and found that *Tnfrsf4*, *Uba52*, *Itgb1*, *mt-Nd4I*, and *Ccl5* were the top 5 most upregulated genes in MASH CD4^+^ T cells (Fig. 6A). Tnfrsf4 (TNF receptor superfamily member 4), encoding the OX40 costimulatory receptor on antigen-experienced T cells, was also specifically expressed by subsets enriched in MASH livers, such as Th1 and Treg cells, including some cytotoxic cells, both in murine and human MASH (Fig.6B, Fig. S6A). We next examined the protein expression of OX40 and its only known ligand, OX40L, in isolated intrahepatic leukocytes from chow and FFC-fed mice by flow cytometry (Fig. S6A-C). Within CD45^+^ intrahepatic leukocytes, OX40 expression was mainly detected in the CD4^+^ T-cell compartment (Fig. 6C). In line with the transcriptomic data, we found that MASH livers have a significantly higher percentage of OX40^+^ CD4^+^ T cells (Figure 6D). OX40 expression was virtually absent in intrahepatic CD8^+^ T cells and non-T leukocytes (around 2% and 0.5%, respectively), confirming that OX40 is mainly associated with hepatic CD4^+^ T cells (Fig. 6D). Given the specific increase of OX40 in hepatic CD4^+^ T cells, we then wonder if it was a liver-specific phenomenon. Evaluating OX40 expression in other tissues during MASH, we found a similar increase in OX40^+^ CD4^+^ T cells in the spleen but not in mesenteric lymph nodes (MLN) or lamina propria (LP) (Fig. 6E). OX40^+^ CD4^+^ T cells were also slightly increased in PBMC, which was consistent with the CD4^+^ T-cell phenotype observed in the blood (Fig. 3). OX40 expression in CD8^+^ T cells in these tissues was low (Fig. 6E). We next examine the expression of OX40 ligand (OX40L), the only known cognate ligand of OX40. OX40L expression was low in liver leukocytes under normal conditions but was markedly increased in leukocytes from MASH livers (Fig. 6F). This increase was mainly driven by an increased proportion of OX40L^+^ macrophages, with a slight increase in OX40L^+^ dendritic cells (Fig. 6G-H). Notably, this striking increase in OX40L leukocytes during FFC feeding was only detected in the liver but not in the spleen, MLN, LP or PBMCs (Fig. 6I). In human MASH, we analyzed OX40L and OX40 hepatic expression using a bulk RNAseq dataset (GEO: GSE126848)^13^ as scRNAseq data for liver leukocytes were not available. Hepatic OX40L transcript levels were increased in MASH patients compared to healthy or obese controls, while hepatic OX40 expression showed an increasing tendency in MASH without reaching statistical significance (Fig. S6E). Collectively, this thorough assessment establishes the OX40L-OX40 axis as a CD4^+^ T cell-specific target for MASH.

**Figure 6.**
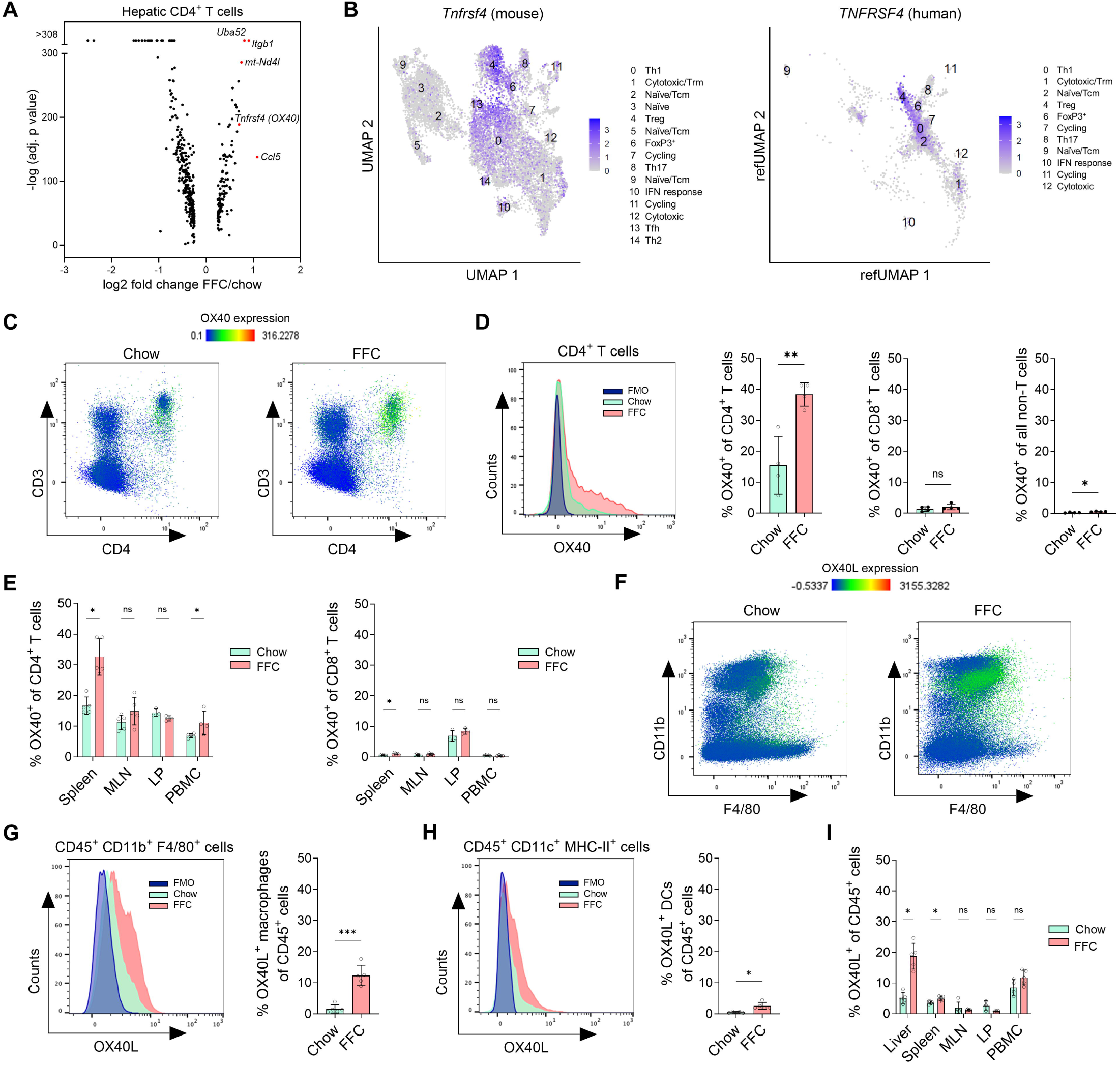
Tnfrsf4/OX40 is upregulated in CD4^+^ T cells in MASH. **(A)** scRNAseq pseudo-bulk differential expression analysis for intrahepatic CD4^+^ T cells from chow and FFC-fed mice. The top five upregulated genes are shown in red. **(B)** Expression of Tnfrsf4/TNFRSF4 (OX40) across intrahepatic CD4^+^ T-cell subsets in chow versus FFC-fed mice and human MASH. **(C)** Representative intensity heatmap of OX40 expression on flow-gated CD45^+^ cells across CD3 and CD4 expression in chow and FFC liver leukocytes. **(D)** Frequency of CD4^+^ T cells, CD8^+^ T cells, and non-T cells expressing OX40 in the liver. **(E)** Frequency of CD4^+^ and CD8^+^ T cells expressing OX40 in the spleen, MLN, LP, and PBMC. **(F)** Representative intensity heatmap of OX40L expression on flow-gated CD45^+^ cells across CD11b and F4/80 expression in chow and FFC liver leukocytes. **(G)** Frequency of macrophages (CD11b^+^ F4/80^+^) expressing OX40L in the liver. **(H)** Frequency of dendritic cells (CD11c^+^ MHC-II^+^) expressing the OX40L in the liver. **(I)** Percentage of leukocytes expressing OX40L in liver, spleen, PBMC, MLN, and LP. N of samples indicated in the graphs. N of samples indicated in the graphs. *p < 0.05, **p < 0.01, ***p < 0.001. ns, non-significant. Abbreviations: Tnfrsf4, TNF Receptor Superfamily Member 4; FMO, Fluorescence Minus One; FFC, high fat, fructose, and cholesterol; PBMC, peripheral blood mononuclear cells; MLN, mesenteric lymph nodes; LP, lamina propria.

### Blocking the OX40L-OX40 axis reverses MASH and hepatic fibrosis in mice

Confirming that OX40 expression is restricted to CD4^+^ T cells in MASH livers, we then aimed to therapeutically inhibit OX40 function in established MASH using a neutralizing monoclonal antibody against OX40L, the only known ligand for OX40. After 26 weeks on the FFC diet, mice were randomized to receive either anti-OX40L antibody (α-OX40L) or control IgG three times a week for four weeks while continuing the FFC diet (Fig. 7A). Treatment with α-OX40L had no effect on body weight, liver weight and white adipose tissue weight (Fig. 7B-C, Fig. S7A-B). Likewise, liver injury and steatosis were not different between α-OX40L and IgG-treated FFC-fed mice (Fig. 7D-E). However, assessing liver histology, it became evident that α-OX40L treatment attenuated disease severity and tissue fibrosis. Liver tissue samples were scored by a blinded pathologist, revealing a significant reduction in both the NAFLD activity score (NAS) and portal inflammation following α-OX40L treatment (Fig. 7F-G, Fig. S7C). To get better insight into the liver immune environment upon blocking the OX40L-OX40 axis, we conducted a comprehensive analysis by mass cytometry (CyTOF) on intrahepatic leukocytes using an antibody panel for annotating major immune cell subsets. This approach identified 25 clusters of hepatic immune cells based on their distinct marker expression (Fig. S7D-E). Furthermore, unsupervised hierarchical clustering based on cluster frequencies revealed that the samples clustered within their respective experimental group, demonstrating a distinct hepatic immune cell composition driven by diet and treatment (Fig. S7F). The most pronounced effect of OX40L-OX40 blockade in FFC-fed mice was represented by a decreased abundance of Ly6C^+^ monocyte-derived macrophages (Fig. S7G), indicating that the OX40L-OX40 axis is relevant to the crosstalk between CD4^+^ T cells and recruited myeloid cells in MASH. Finally, we evaluated the extent of liver fibrosis. Mice treated with the α-OX40L antibody showed an overall reduction in fibrosis scores provided by a blinded pathologist compared to IgG-treated mice (Fig. 7H). These results were consistent with a significant reduction in liver collagen deposition upon α-OX40L antibody treatment, as confirmed by Sirius red staining quantification and hepatic hydroxyproline assay (Fig. 7I-J). Thus, the treatment with neutralizing anti-OX40L antibody was able to attenuate disease severity and reverse liver fibrosis. To test the targeting of the OX40L-OX40 axis in a human preclinical model, we adopted a protocol for inducing MASLD in human precision-cut liver slices.^14^ Human liver slices incubated in GFIPO media (containing glucose, fructose, insulin, palmitate, and oleate) developed steatosis and upregulation of genes involved in lipogenesis and inflammation (Fig S7H-I). Interestingly, OX40 mRNA was also upregulated by GFIPO media (Fig. S7I). To test OX40 blockade in this human MASLD model, liver slices were incubated in GFIPO media in the presence of Rocatinlimab, an antagonistic OX40 monoclonal antibody (α-OX40) currently studied in clinical trials for atopic dermatitis^15^. Anti-OX40 antibody treatment reduced mRNA expression of key CD4^+^ T cell-associated inflammatory markers, such as IFNγ and CXCL10 (Fig. 7K). In summary, these results indicate that activated CD4^+^ T cells promote MASH pathogenesis through the OX40L-OX40 signaling axis, which can be potentially therapeutically targeted.

**Figure 7.**
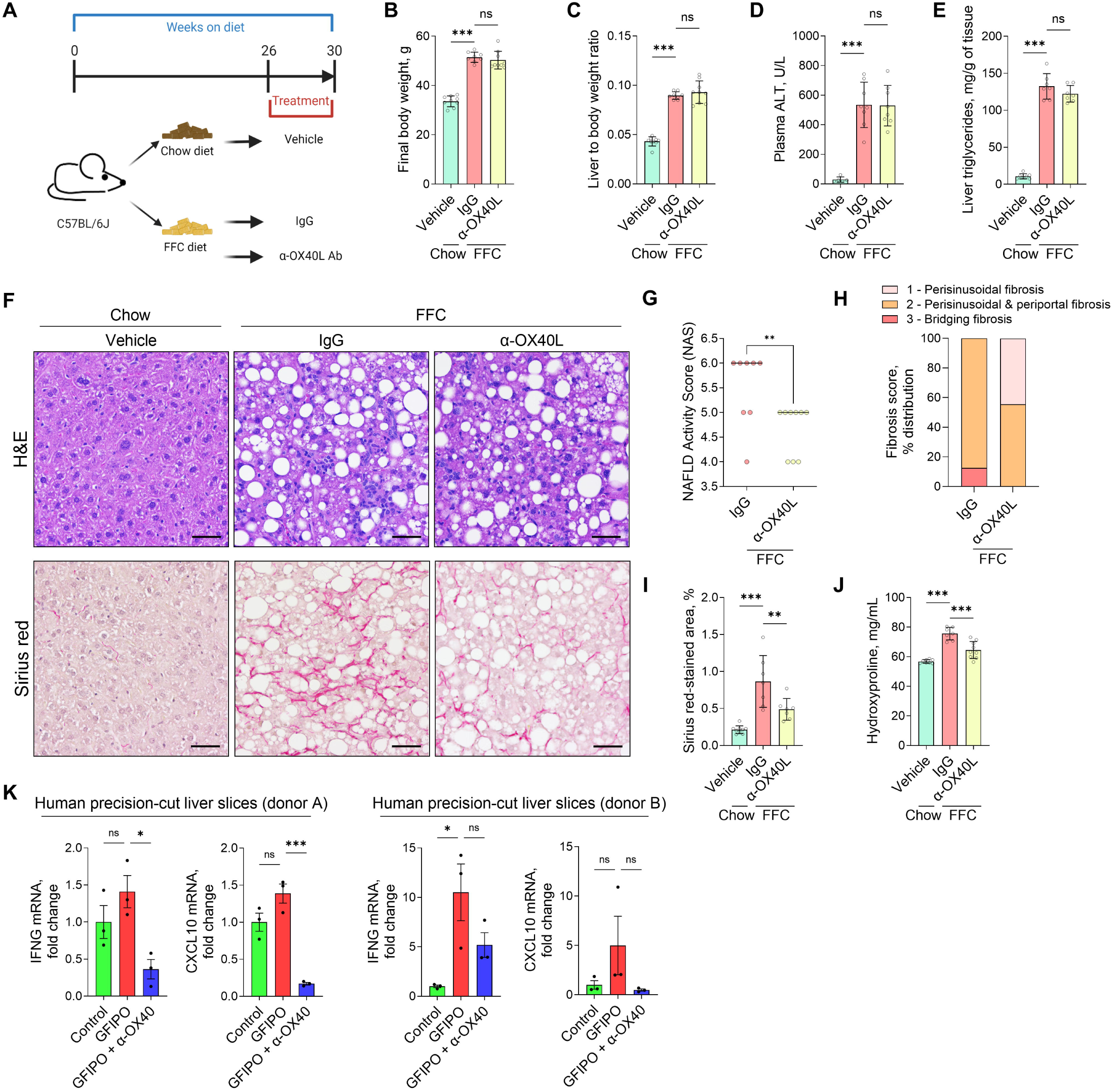
Blocking the OX40L-OX40 axis reverses MASH and hepatic fibrosis in mice. **(A)** Schema of neutralizing monoclonal anti-OX40L antibody treatment. **(B)** Body weight at the time of sacrifice. **(C)** Liver weight normalized to the body weight. **(D)** Liver injury assessed by plasma ALT. **(E)** Hepatic triglyceride quantification. **(F)** H&E and Sirius red staining on liver tissue sections. **(G)** NAFLD activity score (NAS). **(H)** Fibrosis score. **(I)** Sirius red staining quantification. **(J)** Hydroxyproline assay for quantifying liver collagen content. Representative images (taken with a 20× objective) are shown in F; the scale bar is 50 µm. N of samples indicated in the graphs. **p < 0.01, ***p < 0.001. ns, non-significant. Abbreviations: FFC, high fat, fructose, and cholesterol; IgG, immunoglobulin G; Ab, antibody; H&E, Hematoxylin and eosin; ALT, alanine aminotransaminase.

## DISCUSSION

The current studies provide novel insights into the CD4^+^ T-cell landscape in healthy livers and livers affected by MASH. The principal findings from the murine MASH model indicate that: 1) hepatic CD4^+^ T cells are highly heterogeneous, and a MASH-inducing diet skews their polarization toward Th1, cytotoxic and regulatory phenotypes, and 2) pharmacologic inhibition of CD4^+^ T cell costimulatory signals can reverse established MASH and fibrosis. These findings were also validated in human samples and an *ex vivo* model, and are discussed in greater detail below.

Single-cell technologies have enabled a better understanding of the liver immune microenvironment in healthy and MASH livers. Prior studies have reported that various hepatic T cell subsets exist in mouse and human MASH.^16,17^ However, a detailed characterization of the hepatic CD4^+^ T landscape at the protein, transcriptional, and functional levels has remained limited. By utilizing mass cytometry and CITEseq, our study provides a novel single-cell atlas of hepatic CD4^+^ T cells in healthy and MASH livers. Our study also highlights the importance of using protein markers for studying T cells since about 40% of hepatic CD4^+^ T cells had undetectable CD4 mRNA in our scRNAseq experiment. To model experimental MASH, we employed a diet high in fat, fructose, and cholesterol – the FFC diet. The FFC diet, which phenocopies the histological features of MASH and the associated metabolic dysfunction, has been shown to have the highest fidelity and relevance to human MASH.^18,19^ In this model, analyzing the protein expression and transcriptomes of hepatic CD4^+^ T cell populations, we identified CD4^+^ T cell populations in MASH that were enriched for signatures of Th1, Treg, and cytotoxic CD4^+^ T cells, which are novel in the context of MASH. We also found that similar T-cell populations were present in patients with MASH.

IFNγ-producing Th1 cells were the most enriched phenotype in MASH livers, displaying different stages of activation, indicated by varying expression levels of IL7R (memory differentiation) and/or PD-1 (activation or exhaustion marker). *Ex vivo* re-stimulated hepatic CD4^+^ T cells derived from MASH mice showed significantly higher production of IFNγ and TNF-α, but not IL-17A, compared to CD4^+^ T cells derived from healthy mice. This indicates that Th1 cells are both the predominant and primed T-cell population in MASH livers induced by a western diet. Similar trends have been previously reported for hepatic CD4^+^ T cells obtained from patients with MASH.^16^ Th1 cells were also increased in the circulation of our MASH mice, which is consistent with previously reported increases in Th1 or IFNγ-producing CD4^+^ T cells in the peripheral blood of patients with MASH.^20,21^ CD4^+^ regulatory T cells (Tregs), defined as FoxP3^+^ CD25^high^ cells, were also significantly increased in MASH liver, which may indicate that they are being recruited to the inflamed liver as a compensatory mechanism to suppress excessive immune responses and limit tissue damage. However, recent evidence also indicates that Tregs may contribute to disease progression by amphiregulin secretion, which activates a pro-fibrotic transcriptional program in hepatic stellate cells.^22^ Besides Tregs, we also report the presence of a distinct new subset of hepatic FoxP3^+^ with no or low expression of CD25. These FoxP3^+^ CD25^-/low^ cells may represent newly induced Tregs that have not yet fully upregulated CD25, Tregs losing their regulatory phenotype, or cells that transiently upregulated FoxP3 during activation. Based on scRNAseq trajectory analysis, this FoxP3^+^ population seems to be transcriptionally related to both Tregs and Th17 cells. By mass cytometry, we detected two subsets of Th17 cells that were differentiated by the expression of CD103 (integrin αE) and integrin β7. Interestingly, αEβ7 expression is thought to mark proinflammatory colonic CD4^+^ T cells.^23^ Although changes in Th17 populations and IL17 production were relatively minor, it is likely that these cells also contribute to MASH development.^24^

Herein, we describe a novel subset of CD4^+^ T cells exhibiting cytotoxic features that have not been previously reported. First, by mass cytometry, we detected a distinct cluster of CD4^+^ T cells expressing T-bet and Eomes, a transcription factor of cytotoxic lymphocyte lineages. In CITEseq, cluster 1, which corresponds to cytotoxic CD4^+^ T cells expressing T-bet and Eomes, also expressed cytotoxic mediators such as granzymes, perforin, Ifng, and Fas ligand. Cytotoxic CD4^+^ T cells have been described in chronic viral infection, autoimmune diseases, and chronic inflammatory diseases, such as inflammatory bowel disease.^25^ Thus, their presence in MASH suggests that CD4^+^ T cells in MASH are exposed to chronic immune stimulation. Besides cytotoxic CD4^+^ T cells, other novel subsets described herein were CD4^+^ T cells with IFN response and follicular helper cells (Tfh). IFN response cluster expressed genes are typically induced by type I interferon, i.e., IFNα and IFNβ. Tfh cells, which provide help to B cells in secondary lymphoid organs, are not expected to be present in liver tissue. Although Tfh-like cells can be present in the circulation, hepatic Tfh cells described herein by mass cytometry and CITEseq seem to resemble tissue Tfh cells more closely than circulating Tfh-like cells, mainly due to high expression of BCL6 and CD69. Altogether, these protein and RNA single-cell datasets provide a useful resource for a relevant preclinical MASH model commonly used to study disease pathogenesis and evaluate therapeutic approaches.

In terms of CD4^+^ T cell function during MASH, we demonstrated that after 24 weeks on the FFC diet, mice lacking the CD4 gene had a significantly reduced liver injury, inflammation, and fibrosis, compared with the control group, indicating that the absence of CD4^+^ T cells is protective against disease development. Some limitations of this study relate to the global deletion of the CD4 gene, which can lead to compensatory changes in CD8^+^ T cells. For example, the CD4^+^ T cell deficiency is compensated for by developing MHC-II-restricted CD8^+^ T cells or double-negative T cells.^26^ Loss of CD4 may also affect the development of iNKT, albeit the numbers of hepatic iNKT cells (recognizing PBS-57 loaded CD1d tetramer) were not affected in our CD4 knockout mice (not shown). Nevertheless, our results obtained in CD4 knockout mice are consistent with those shown in a humanized mouse model engrafted with human immune cells and fed with a high-fat, high-carbohydrate diet.^27^ In this model, depletion of human CD4^+^ T cells before and during the diet feeding reduced NAS score, plasma proinflammatory cytokines, and the extent of fibrosis. Similarly, inhibition of α4β7^+^ CD4^+^ T cells attenuated liver inflammation and fibrosis in another model utilizing western diet-fed F11r^-/-^ mice, which lack junctional adhesion molecule A and have increased intestinal permeability.^28^ On the contrary, it has been previously reported that steatohepatitis induced by a methionine-choline-deficient diet causes selective loss of CD4^+^ T cells, which drives disease progression.^29^ These conflicting findings are likely explained by the distinct mechanisms by which a methionine-choline-deficient diet and a western-like diet induce steatohepatitis in mice.

In addition to a preventive approach for MASH, we also tested a therapeutic strategy targeting CD4^+^ T cells in already established MASH. Our study identified TNFRSF4/OX40 as one of the most upregulated molecules associated with hepatic Th1 cells. OX40, a costimulatory receptor upregulated upon T-cell activation, has been described to promote cell proliferation, survival, and effector function.^30^ We found that OX40 is selectively upregulated on intrahepatic CD4^+^ T cells, but not CD8^+^ T cells, in MASH and that therapeutic blockade of the OX40L-OX40 axis significantly reduces liver inflammation associated with monocyte-derived macrophages and fibrosis, without affecting steatosis or body weight. This is consistent with emerging evidence that crosstalk between multiple immune cell types collectively contributes to liver inflammation and fibrogenesis.^7^ Moreover, our findings in human precision-cut liver slices provide translational support by demonstrating that OX40 inhibition using Rocatinlimab, an antagonistic antibody investigated in clinical trials,^15^ suppresses inflammatory gene expression during nutrient excess and lipo/glucotoxic stress. These results collectively suggest that targeting the OX40L- OX40 axis could represent a novel, immunologically focused strategy to treat fibrotic progression in MASLD, addressing a current gap in therapeutic options that primarily target metabolic pathways.

## Abbreviations

αSMA,: Alpha-smooth muscle actin;
Ab,: Antibody;
ADT,: Antibody-derived tag;
ALT,: Alanine aminotransferase;
AST,: Aspartate aminotransferase;
CD,: Cluster of differentiation;
CLAMS,: Comprehensive Lab Monitoring System;
CXCL,: C-X-C motif chemokine ligand;
CXCR,: C-X-C motif chemokine receptor;
CyTOF,: Cytometry by Time-of-Flight;
Eomes,: Eomesodermin;
eWAT,: Epidydimal white adipose tissue;
F4/80,: Adhesion G protein-coupled receptor E1;
FABP5,: Fatty acid binding protein 5;
FACS,: Fluorescence-activated cell sorting;
FFC,: High fat, fructose, and cholesterol;
FoxP3,: Forkhead box P3;
GATA3,: Trans-acting T-cell-specific transcription factor;
GFIPO,: Glucose, fructose, insulin, palmitate and oleate;
GPNMB,: Glycoprotein non-metastatic melanoma protein B;
H&E,: Hematoxilin and eosin;
HOMA-IR,: Homeostasis model assessment of insulin resistance;
HPF,: High power field;
ICOS,: Inducible T-cell costimulator;
IFNγ,: Interferon gamma;
IgG,: Immunoglobulin G;
IL,: interleukin;
IL7R,: Interleukin 7 receptor;
iNKT,: Invariant natural killer T;
Itg,: Integrin;
Lgals3,: galectin-3;
LP,: Lamina propria;
Ly6C,: Lymphocyte antigen 6 complex, locus C;
Ly6G,: lymphocyte antigen 6 complex, locus G;
MASH,: Metabolic dysfunction-associated steatohepatitis;
MASLD,: Metabolic dysfunction-associated steatotic liver disease;
MHC-II,: Major histocompatibility complex class II;
MLN,: Mesenteric lymph nodes;
NAFLD,: Nonalcoholic Fatty Liver Disease;
NAS score,: Nonalcoholic fatty liver disease Activity Score;
OX40,: Tumor necrosis factor receptor superfamily member 4;
OX40L,: Tumor necrosis factor superfamily member 4;
PBMC,: Peripheral blood mononuclear cell;
PCLS,: Precision-cut liver slice;
PD1,: Programmed cell death protein 1;
PMA,: Phorbol 12-myristate 13-acetate;
qPCR,: Quantitative polymerase chain reaction;
RNAseq,: RNA sequencing;
RORγT,: RAR-related orphan receptor gamma T;
scRNAseq,: single- cell RNA sequencing;
SPP1,: Secreted phosphoprotein 1/osteopontin;
T-bet,: T-box transcription factor TBX21;
Tcm,: central memory T cell;
TCRβ,: T-cell receptor beta;
Tfh,: T follicular helper;
TGFβ,: Transforming growth factor beta;
Th,: T helper;
TNF,: Tumor necrosis factor;
Treg,: Regulatory T cell;
TREM2,: Triggering receptor expressed on myeloid cells 2;
tSNE,: t-distributed stochastic neighbor embedding;
UMAP,: Uniform manifold approximation and projection;
WT,: Wild-type.

## Acknowledgments

The authors thank the staff of the Mayo Clinic Immune Monitoring Core, Samera Farwana and Kaitlyn Whitaker, for their assistance with mass cytometry and PhenoCycler experiments. The authors appreciate the support from the Clinical Core and Optical Microscopy Core of the Mayo Clinic Center for Cell Signaling.

## Lead Contact

Further information and requests for resources and reagents should be directed to and will be fulfilled by the lead contact, Petra Hirsova.

## Materials Availability

This study did not generate new unique reagents.

