## Supplemental Figures and Methods for "CD4^+^ T cells promote fibrosis during metabolic dysfunction-associated steatohepatitis"

**The PDF file includes:**

Materials and Methods
Figs. S1 to S7
Tables S1 to S7
References

#### Materials and Methods

##### Animal studies and experimental design

All animal studies were performed at Mayo Clinic and in accordance with and approved by the Institutional Animal Care and Use Committee. C57Bl6/J (#000654) and $CD4^{-/-}$ (#002663) mice were purchased from the Jackson Laboratory (Bar Harbor, ME, USA). All mice used had a C57Bl/6J background. Mice were co-housed in cages using corn cob bedding. Mice were kept at the Mayo Clinic Rochester animal facility with a 12-hour light-dark cycle. All mice were placed on an experimental diet at 10 weeks of age. The chow diet was a standard rodent diet (PicoLab 5053, LabDiet) with tap water. FFC diet contains high fat (40% calories) and high cholesterol (0.2%) (#AIN-76A Western Diet, TestDiet), with fructose (18.9 g/L) and glucose (23.1 g/L) added to the drinking water (total sugar 42 g/L) as previously described.[1-4] Male C57Bl/6J and $CD4^{-/-}$ mice were fed chow or FFC diet for 24 weeks. For the pharmacologic treatment study, male C57Bl/6J were on chow or FFC diet for 26 weeks, after which they were randomized to receive 300 µg of anti-OX40L antibody (#BE0033-1, BioXCell) or control IgG (#BE0090, Bio X Cell) intraperitoneally (i.p.) three times a week for four weeks while continuing the diets. The antibodies were diluted in pH 7.0 Dilution Buffer (#IP0070, BioXCell). For hepatic or circulating $CD4^+$ T cell characterization experiments (e.g., mass cytometry, CITEseq, ex vivo culture), male C57Bl/6J mice were fed chow or FFC diet for 24 weeks. At the end of the feeding period, mice were sacrificed under general anesthesia induced by a combination of xylazine and ketamine. The blood, epididymal fat, spleen, mesenteric lymph nodes, lamina propria, and liver were harvested and processed for further examination.

##### Murine primary cell isolation

Hepatic leukocytes were isolated using the liver dissociation kit (#130-105-807, Miltenyi Biotec) and Percoll (#17-0891-01, GE Healthcare) gradient as we described previously.[4-6] Hepatic leukocytes were subjected to flow cytometry or mass cytometry by time-of-flight. For the analysis of CD4 T cells, mouse livers were flushed to remove circulating lymphocytes, followed by liver leukocyte isolation as described above and subsequent enrichment of CD4 T cells via positive selection using magnetic beads separation (#130-117-043, Miltenyi Biotec). For the analysis of peripheral blood mononuclear cells (PBMCs), fresh mouse blood was collected in lithium heparin tubes and subsequently subjected to a Ficoll-Paque (#17-1440-02, GE Healthcare) gradient according to the manufacturer's instructions. For the analysis of splenocytes, the spleen was harvested and mechanically dissociated by gently crushing between two frosted glass slides, filtered using a 70 µm cell strainer, and treated with a Red Blood Cell Lysis Solution (#130-094-183, Miltenyi Biotec) to remove erythrocytes. To achieve a single-cell suspension, splenocytes were subsequently filtered using a 40 µm cell strainer. For the analysis of mesenteric lymph nodes (MLNs), MLNs were harvested by gently removing the mesentery and fat, followed by mechanical dissociation through a 70 µm cell strainer using the plunger of a syringe. For the analysis of lamina propria (LP), the intestine was collected from mice and prepped as described previously by us.[7]

##### Mass cytometry by time-of-flight

High-dimensional mass cytometry by time-of-flight (CyTOF) was used to examine the immune composition of mouse hepatic CD4 T cells, mouse PBMCs, and mouse hepatic leukocytes. Cells were stained with a custom panel of heavy-metal conjugated antibodies (Table S1-S3). Data normalization, cleanup, and analysis were performed by the Mayo Clinic Immune Monitoring Core as previously described by us.[4-6] Data in .fcs files were normalized using CyTOF Software (version 6.7.1014). For analysis of $CD4^+$ T cells from liver tissue or PBMCs, we gated on $CD3^+$ $TCR\beta^+$ $CD4^+$ $CD8^-$ $NK1.1^-$ cells to perform the downstream analysis. Cleaned .fcs files were analyzed by the R-based tool Cytofkit version 3.8. The R-phenograph algorithm clustering and dimensionality reduction were used for all markers in the panel. Clusters were visually presented as tSNE maps and heatmaps.

**Metabolic phenotyping**

The extent of insulin resistance was assessed at 22 weeks of feeding for C57Bl/6J and $CD4^{-/-}$ mice. Mice were fasted overnight for 12 hours in clean cages without access to food, and sugar water was replaced with regular water. Fasting glucose was measured by a blood glucose meter (#560001, Assure Platinum) and test strips (#500050, Assure Platinum) from the tail nick blood. Plasma insulin concentration was determined using an ultrasensitive mouse insulin enzyme-linked immunosorbent assay kit (#90080, Crystal Chem) following the manufacturer's instructions. Fasting glucose and fasting insulin values were used to calculate the homeostasis model assessment of the insulin resistance (HOMA-IR) index as described.[3,4] At 20 weeks of feeding, mice on chow and FFC diet were placed in fully automated metabolic cages known as Comprehensive Lab Monitoring System or CLAMS (Columbus Instruments) as described previously by us. The system monitors metabolic parameters by indirect calorimetry, feeding behavior, and voluntary activity. Where indicated, indirect calorimetry data from CLAMS were normalized to lean body mass. Body composition (fat mass and lean mass) was measured with EchoMRI-100 analyzer (EchoMRI LLC) as described previously by us.[3,4]

**Biochemical analysis**

Plasma alanine aminotransferase (ATL) levels were measured using a veterinary chemistry analyzer VS2 (#500-0040-12, VetScan) using the Mammalian Liver Profile (#10023223, VetScan). Plasma aspartate transaminase (AST) levels were measured using the AST Assay Kit (#MAK055, Sigma-Aldrich) as per the manufacturer's instructions. Liver triglyceride levels were measured in the mouse liver homogenates using the EnzyChrome triglyceride assay kit (#ETGA200, BioAssay Systems) as described before.[3,4] Liver hydroxyproline levels as a direct measure of collagen content were determined using frozen liver specimens as described before.[8]

**Histopathology**

The harvested mouse livers were diced into 5 mm x 5 mm pieces, fixed in 10% neutral buffered formalin for 24 hours, and embedded in paraffin. Tissue sections (5 µm) were cut using a microtome (Reichert Scientific Instruments) and positioned on glass slides to achieve formalin-fixed paraffin-embedded (FFPE) sections. Hematoxylin and eosin (H&E) staining was performed using standard techniques in Mayo Clinic's Histology Core. Mouse FFPE liver tissue sections were stained with Picrosirius Red or the Sirius Red/Fast Green Collagen Staining Kit (#9046, Chrondrex) for liver fibrosis assessment. Random fields (10-20 per sample) were quantified by polarized light using Nikon Eclipse TE300 with 10× objective in a blinded manner. The data are shown as a percentage of Sirius red-positive area. The non-alcoholic fatty liver disease activity score (NAS), a quantitative score that assesses steatosis, ballooned hepatocytes, and lobular inflammation, and fibrosis score (using Sirius Red-stained slides) was determined by a blinded liver pathologist. Representative images were taken (Scope A1, Zeiss) using 20× objective.

**Immunohistochemistry**

FFPE mouse liver tissue sections were deparaffinized, hydrated, and incubated with specific antibodies using the following dilutions: Lgals3 1:250 (#14-5301-82, eBioscience), αSMA 1:1000 (#ab12496, Abcam), F4/80 1:500 (#70076, Cell Signaling Technology) for 1 h at room temperature or overnight at 4°C. Bound antibodies were detected using a Vectastain ABC kit (#PK-4001, #PK-6102, Vector Laboratories), diaminobenzidine as a substrate, and counterstaining with hematoxylin. Positive staining in 20 high-power random fields was quantified using Nikon Eclipse TE300 microscope software in a blinded manner. Representative images were taken (Scope A1, Zeiss) using 20× objective.

***Ex vivo* experiments with murine CD4 T cells**

Purified hepatic $CD4^+$ T cells were cultured in anti-CD3 (#553057, BD Biosciences) pre-coated plates in a complete RPMI-1640 (#11875093, Gibco) medium supplemented with 10% FCS, 10 mmol/L HEPES pH 7.4, 1 mmol/L sodium pyruvate, 2 mmol/L L-glutamine, 1% non-essential amino acids, penicillin, streptomycin, and 50 µmol/L β-mercaptoethanol, and anti-CD28 (#553294, BD Biosciences, 2 µg/ml).[7] Cells were subsequently stimulated with PMA/Ionomycin (#423302, BioLegend, 1:500 dilution) or vehicle (DMSO) for 6 hours in the presence of Brefeldin A (#420601, BioLegend, 1:1000 dilution) for flow cytometric analysis of intracellular cytokines or without Brefeldin A for measurement of cytokines in cell culture supernatants. Following stimulation, cells treated with Brefeldin A were harvested, washed in 1X PBS, fixed using 4% paraformaldehyde, and subjected to flow cytometric analysis. Supernatants from cells stimulated without Brefeldin A were used for bead-based multiplex cytokine assay using LEGENDplex™ Mouse Th Cytokine Panel (#741044 , BioLegend) according to the manufacturer’s instructions.

**Flow cytometry**

Single-cell suspensions were obtained before all flow cytometric analyses. Briefly, cells were incubated with TruStain FcX PLUS (anti-mouse CD16/32) Antibody (#156603, BioLegend) and Viobility Fixable Dyes (#130-130-404 or 130-130-403, Miltenyi Biotec) where indicated, for 10 minutes at 4°C in the dark. Cells were then washed and labeled with fluorochrome-conjugated cell surface antibodies (Table S4) for 20 minutes at 4°C in the dark. For intracellular markers, cells were subsequently washed, fixed, and permeabilized with Cyto-Fast™ Fix/Perm Buffer Set (#426803, BioLegend) and labeled with fluorochrome-conjugated intracellular antibodies for 20 minutes at room temperature in the dark. Flow cytometry was performed with the MACSQuant Analyzer 10 (Miltenyi Biotec) or the LRSFortessa X20 (BD Biosciences). Data were analyzed and visualized in FlowJo software V10.10 (BD Life Sciences). Gates were established using fluorescence-minus-one (FMO) controls.

**Flow cytometric sorting of liver monocyte-derived macrophages**

Hepatic leukocytes were stained with an Fc block (#130-092-575, Miltenyi), viability dye (VioBlue, #130-130-420, Miltenyi) and fluorophore-conjugated antibodies against CD45, CD11b, F4/80, Ly6G (Supplemental Table 4). Monocyte-derived macrophages were defined as $CD45^+$ $CD11b^+$ $F4/80^+$ $Ly6G^-$ cells. Flow sorting was performed on a BD FACSMelody Cell Sorter instrument, and cells were collected into FBS-containing buffer. Cells were then lyzed in TRIzol Reagent (#15596018, Invitrogen) for mRNA analysis by qPCR.

**NanoString nCounter assay**

The expression of 770 genes in murine hepatic $CD4^+$ T cells was assessed by a NanoString nCounter system and mouse Autoimmune Profiling Panel (#115000269, NanoString Technologies). Mouse livers from chow (n=3) and FFC-fed (n=3) mice were flushed to remove circulating lymphocytes, and $CD4^+$ T cells were isolated from hepatic leukocytes using magnetic bead separation (Miltenyi Biotec, #130-117-043) and lyzed in TRIzol Reagent (#15596018, Invitrogen). Total RNA was isolated using Direct-zol RNA Microprep Kits (#R2060, Zymo Research). All following procedures, including sample preparation (100 ng of RNA per sample), hybridization (at 65°C for 17 h), detection, and subsequent analysis, were performed according to the manufacturer's instructions. Data quality control and background thresholding were performed on nSolver using built-in negative controls and housekeeping genes.

**Cellular Indexing of Transcriptomes and Epitopes by Sequencing (CITE-seq ) analysis of hepatic $CD4^+$ T cells**

Mouse livers were flushed with PBS prior to harvesting to remove circulating lymphocytes. Intrahepatic leukocytes were isolated, and $CD4^+$ T cells were enriched using magnetic bead separation (#130-117-043, Miltenyi Biotec). Cells were first incubated with Fc block (#156603, BioLegend) and viability dye (Viobility, Miltenyi Biotec), then stained with a TotalSeq-B CITE-seq (BioLegend) antibody cocktail (Table S5), along with fluorophore-conjugated anti-CD3ε (#152322, BioLegend) and anti-CD4 (#100421, BioLegend) antibodies. Viable $CD3ε^+CD4^+$ cells were purified by fluorescence-activated cell sorting (FACS) using the BD FACSMelody Cell Sorter instrument. Single-cell cDNA libraries were prepared using the 10x Genomics Chromium Single Cell 3' Reagent Kit v2, and sequencing was performed on an Illumina NextSeq 2000 platform at the Mayo Clinic Genome Analysis Core. The sequencing depth was ~100,000 reads per cell for gene expression (GEX) and ~25,000 reads per cell for CITE-seq. Raw data were processed using 10x Genomics Cell Ranger software (v3.0). Downstream analyses were conducted in R (v4.1.0) using Seurat (v3.1.1). Only RNA expression data were used for dimensionality reduction and clustering. Antibody-derived tag (ADT) data were excluded from clustering and were used only for downstream annotation and validation of cell types. Dimensionality reduction and visualization of cell clusters were performed using Uniform Manifold Approximation and Projection (UMAP). Differential gene expression was assessed using the Wilcoxon rank-sum test (default setting in Seurat). Developmental trajectories of $CD4^+$ T cells were inferred using the Slingshot package. GSEA was performed using the fgsea R package (v1.18.0) on genes ranked by average log fold change from Seurat's FindMarkers output. Gene sets were obtained from MSigDB (v7.0). To compare human and mouse $CD4^+$ T cells, human scRNA-seq data (GEO accession: GSE217235) were projected onto our mouse reference using Seurat (v3.1.1). One-to-one orthologous genes were identified using Ensembl BioMart and used to align the datasets. Both datasets were independently normalized and processed for variable feature selection. Integration was performed using Seurat's FindTransferAnchors and MapQuery functions, with the mouse dataset as the reference. Projected human cells were visualized in the mouse UMAP space, and predicted cluster identities were transferred based on nearest neighbors.

**Multiplexed imaging of human liver samples using PhenoCycler (CODEX)**

All human specimens were obtained under protocols approved by the Mayo Clinic Institutional Review Board, and all subjects gave written informed consent. De-identified human liver specimens from patients with MASL (i.e., isolated steatosis) and MASH (stage F0-1) were obtained. The MASL cohort consisted of 4 patients (2 males, 2 females) of age 51 ± 10 years (mean ± SD) and body mass index of 45 ± 5 kg/m2 (mean ± SD). The MASH patient group consisted of 4 patients (2 males, 2 females) of age 48 ± 19 years (mean ± SD) and body mass index of 45 ± 9 kg/m2 (mean ± SD). Both groups of patients underwent liver biopsy or surgical hepatic resection at Mayo Clinic, Rochester, MN, USA. The diagnosis of MASH was based on histologic categorization according to established MASH criteria as assessed by an experienced hepatopathologist. Patients with steatohepatitis caused by drug-induced liver disease and patients with other chronic liver diseases (e.g., cholestatic liver disease, hemochromatosis, excessive alcohol consumption, viral hepatitis, Wilson disease, and alpha-1-antitrypsin deficiency) were excluded. Formalin-fixed paraffin-embedded de-identified human liver samples of isolated steatosis and MASH F0-1 were subjected to high-dimensional spatial imaging using the Phenocycler™ platform (Akoya Biosciences), formerly known as CODEX. Tissue sections (5 µm thick) were mounted on glass coverslips, deparaffinized, rehydrated, and subjected to antigen retrieval following the manufacturer's protocol. A custom antibody panel (Table S6) consisting of DNA-barcoded antibodies targeting immune, stromal, and parenchymal markers was applied to each section. Conjugated antibodies were validated and titrated prior to multiplexing. Imaging was performed using iterative oligonucleotide hybridization, imaging, and signal removal cycles on a Keyence BZ-X800 microscope equipped for automated Phenocycler workflows. Raw image data were processed using Akoya's CODEX Processor and Phenochart software for stitching, background correction, and cell segmentation.

**Human Precision-Cut Liver Slice (PCLS) preparation**

All human specimens were obtained under protocols approved by the Mayo Clinic Institutional Review Board. Liver tissue was collected from consented patients undergoing liver resections. Following resection, tissue was immediately transported in RPMI medium (#11875085, Gibco) at 4 °C and used to minimize ischemic injury. The liver capsule and any cauterized scar tissue were carefully removed to isolate the fresh tissue. Tissue was then sectioned into ~8 mm × 8 mm wedges and embedded in 2.5% low-melting-point agarose (#VF-AGT-VM, Precisionary). Embedded tissue was sliced into 250 µm-thick sections using a Compresstome instrument (#VF-510-0Z, Precisionary), following the manufacturer's recommended blade oscillation and speed settings. Slices were temporarily stored in cold HBSS on ice until sample processing was finished. PCLS samples were then transferred to sterile culture media consisting of a 1:1 mixture of Williams' E Medium (#12551032, Gibco) and

DMEM (#11995-065, Gibco), supplemented with 10% fetal bovine serum (#s11150H, R&D Systems), 1% penicillin-streptomycin (#15140122, Gibco), and Insulin-Transferrin-Selenium-Ethanolamine (ITS-X, #51500056, Gibco). To mitigate stress-induced signaling from the slicing process, the media was further supplemented with 25 mM valproic acid (#P4543, Sigma-Aldrich), 94 nM SB431542 (#1614, Tocris Bioscience), and 6.7 nM CHIR99021 (#4423, Tocris Bioscience) for the first 24 hours. After that, according to a previously published protocol,[9] slices were incubated for an additional 96 hours in control media or GFIPO media with either Rocatinlimab, an anti-human OX40 antagonistic antibody (10 μg/ml, # HY-P99955, MedChemExpress) or control IgG (10 μg/ml, # HY-P99001, MedChemExpress). The media were refreshed every 24 hours. Slices were cultured at 37 °C with 5% $CO_2$ on a rocking platform (90 rpm) to ensure adequate nutrient and oxygen exchange. Slices were harvested for RNA isolation (TRIzol reagent, #15596018, Invitrogen) or fixed in formalin for histology. Total RNA was isolated using Direct-zol RNA Microprep Kits (#R2060, Zymo Research).

**Gene expression by quantitative real-time polymerase chain reaction (qPCR)**

Liver tissue or cells were homogenized using TRIzol reagent (#15596018, Invitrogen), and the total RNA was isolated (#R2070, Zymo Research). The iScript cDNA Synthesis Kit (#1708891, Bio-Rad) was used to transcribe RNA to cDNA following the manufacturer's instructions. The mRNA levels were quantified by real-time qPCR using Power SYBR Green PCR Master Mix (#4367659, Applied Biosystems) on a QuantStudio 6 Flex instrument (Applied Biosystems). The primers are listed in Table S7. Target gene expression was calculated using ΔΔCt method. Expression of 18S, which was stable across experimental groups, was used for the normalization. Data are shown as a fold change over the expression levels in the control condition.

**Statistical analysis**

Data are presented as means ± SEM. Differences among three or more groups were analyzed using one-way analysis of variance (ANOVA) followed by the Tukey post hoc test. A two-tailed unpaired t-test was used to detect differences between two groups. A P value < 0.05 was considered statistically significant. All analyses were performed using GraphPad Prism 10.0 software.

### Supplemental Figures
#### Figure S1

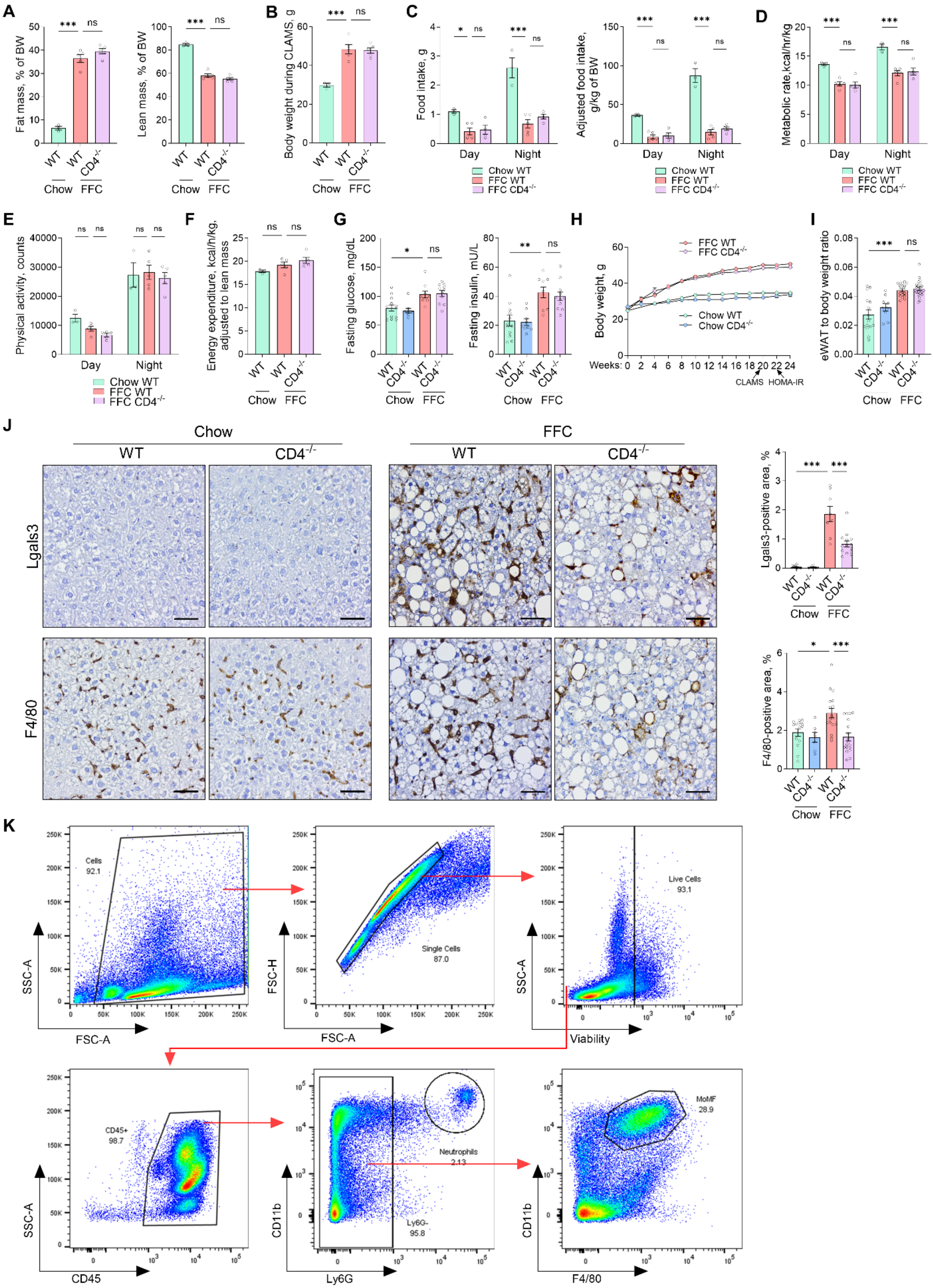

**Fig. S1. CD4 deletion does not affect metabolic phenotype but decreases macrophage-associated inflammation in FFC diet-fed mice. (A-F)** Metabolic parameters were evaluated after 20 weeks of feeding. (A) Body composition measured by EchoMRI. **(B)** Mouse body weight during housing in Comprehensive Lab Animal Monitoring System (CLAMS). **(C-F)** Parameters measured by CLAMS: food intake (C), metabolic rate (D), physical activity (E), and energy expenditure (F). **(G)** Fasting glucose and fasting insulin in the blood were assessed after 22 weeks of feeding. **(H)** Mice were weighed every other week to monitor weight changes. **(I)** Epidydimal white adipose tissue (eWAT) weight normalized to body weight at the time of sacrifice. **(J)** Immunohistochemistry for macrophage markers Lgals3 and F4/80. The positive stain was quantified in a blinded manner. Representative images are shown; the scale bar is 50 μm. **(K)** Gating strategy for purifying hepatic monocyte-derived macrophages by flow sorting. N of samples indicated in the graphs. *p < 0.05, **p < 0.01, ***p < 0.001, ns, non-significant. Abbreviations: CLAMS, Comprehensive Lab Animal Monitor System; eWAT, epidydimal white adipose tissue.

**Figure S2**

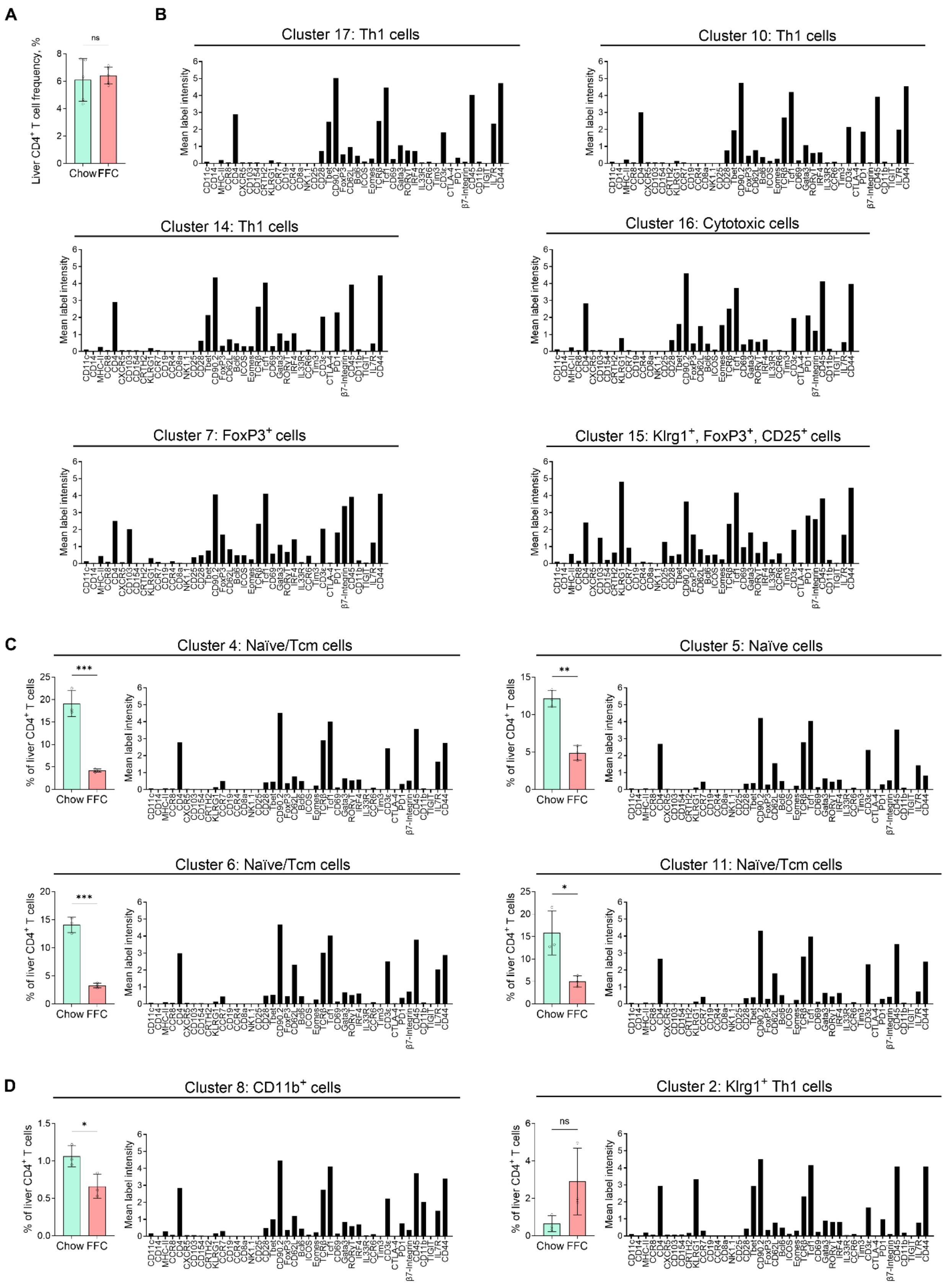
A
B
Cluster 17: Th1 cells
Cluster 10: Th1 cells
Cluster 14: Th1 cells
Cluster 16: Cytotoxic cells
Cluster 7: FoxP3+ cells
Cluster 15: Klrg1+, FoxP3+, CD25+ cells
C
Cluster 4: Naïve/Tcm cells
Cluster 5: Naïve cells
Cluster 6: Naïve/Tcm cells
Cluster 11: Naïve/Tcm cells
D
Cluster 8: CD11b+ cells
Cluster 2: Klrg1+ Th1 cells
Liver CD4+ T cell frequency, %
% of liver CD4+ T cells
Mean label intensity
Chow
FFC
ns
***
**
*

**D (cont.)**

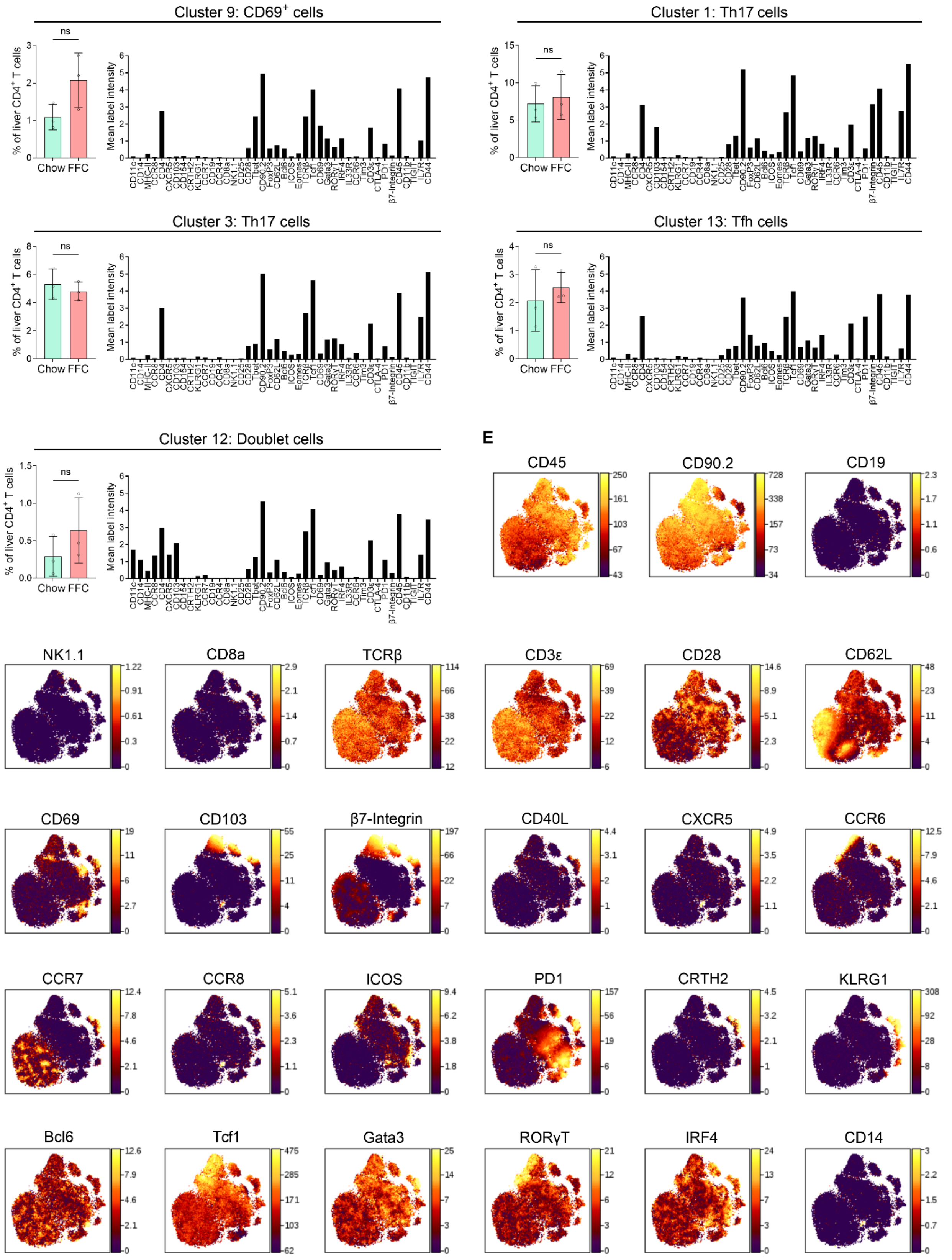


**Fig. S2. Chow livers contain CD4⁺ T cells with naïve-like phenotype. (A)** Frequencies of CD4⁺ T cells within intrahepatic leukocytes (CD45⁺ cells). **(B)** The mean label intensity for the Th1, cytotoxic CD4⁺, and Treg cell clusters that are shown in Fig. 1 G-I. **(C)** Relative abundance and mean label intensity for naïve/central memory CD4 T cell clusters. (D) Relative abundance and mean label intensity for remaining cell clusters. **(E)**

Representative heat maps of protein markers. Abbreviations: FFC, high fat, fructose, and cholesterol; Tfh, T follicular helper cells; *p < 0.05, **p < 0.01, ***p < 0.001, ns, non-significant.

### Figure S3

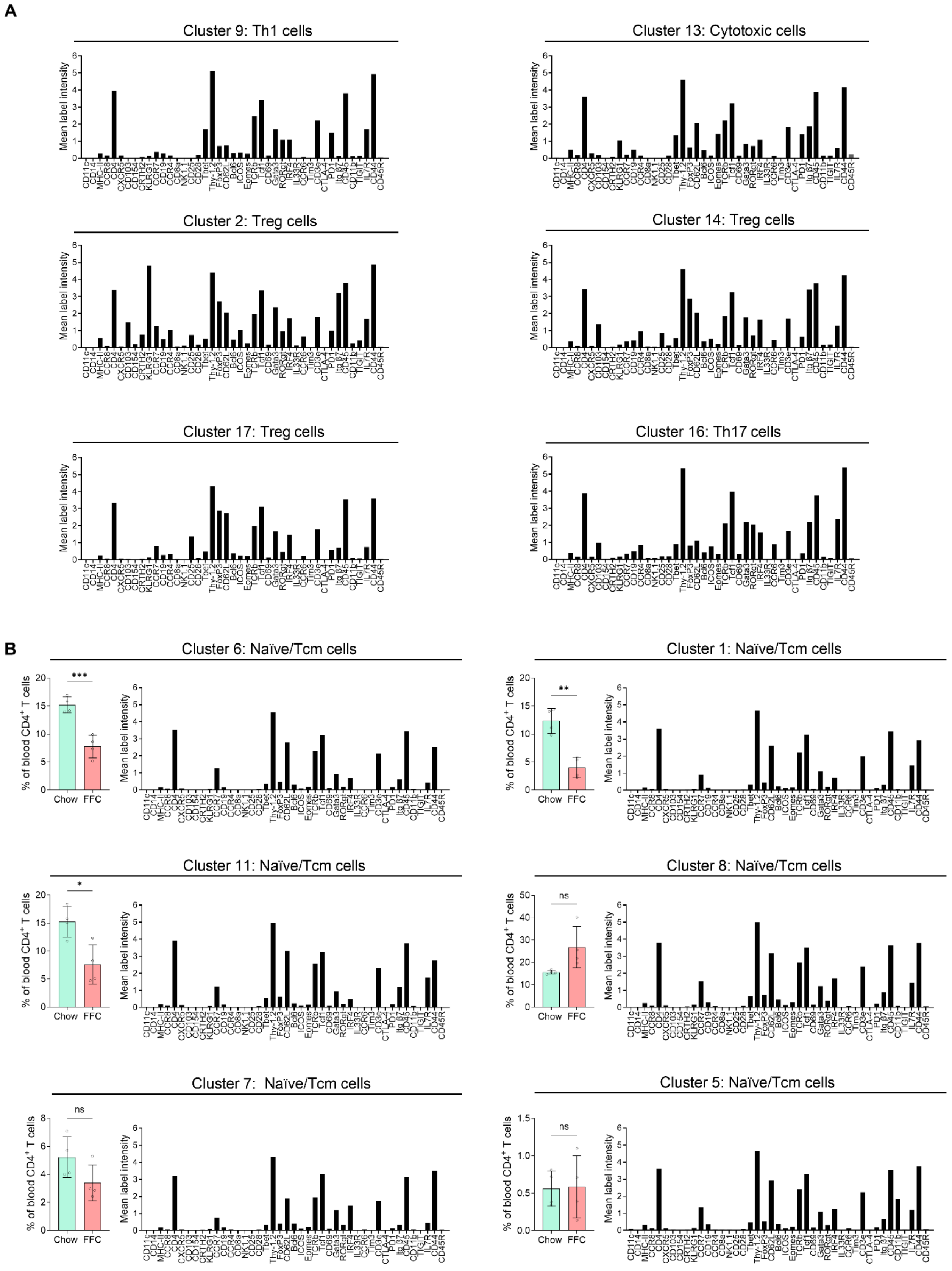
A
Cluster 9: Th1 cells
Cluster 13: Cytotoxic cells
Cluster 2: Treg cells
Cluster 14: Treg cells
Cluster 17: Treg cells
Cluster 16: Th17 cells
Mean label intensity
CD11c CD14 MHC-II CCR8 CD4 CXCR5 CD103 CD154 CRTH2 KLRG1 CCR7 CD19 CCR4 CD8a NK1.1 CD25 CD28 Tbet Thy-1.2 Foxp3 CD62L Bcl6 ICOS Eomes TCRb Tcf1 CD69 Gata3 RORgt IRF4 IL33R CCR6 Tim3 CD3e CTLA-4 PD1 Itg β7 CD45 CD11b TIGIT IL7R CD44 CD45R
B
Cluster 6: Naïve/Tcm cells
Cluster 1: Naïve/Tcm cells
Cluster 11: Naïve/Tcm cells
Cluster 8: Naïve/Tcm cells
Cluster 7: Naïve/Tcm cells
Cluster 5: Naïve/Tcm cells
% of blood CD4+ T cells
Chow
FFC
***
**
*
ns

**B (cont.)**

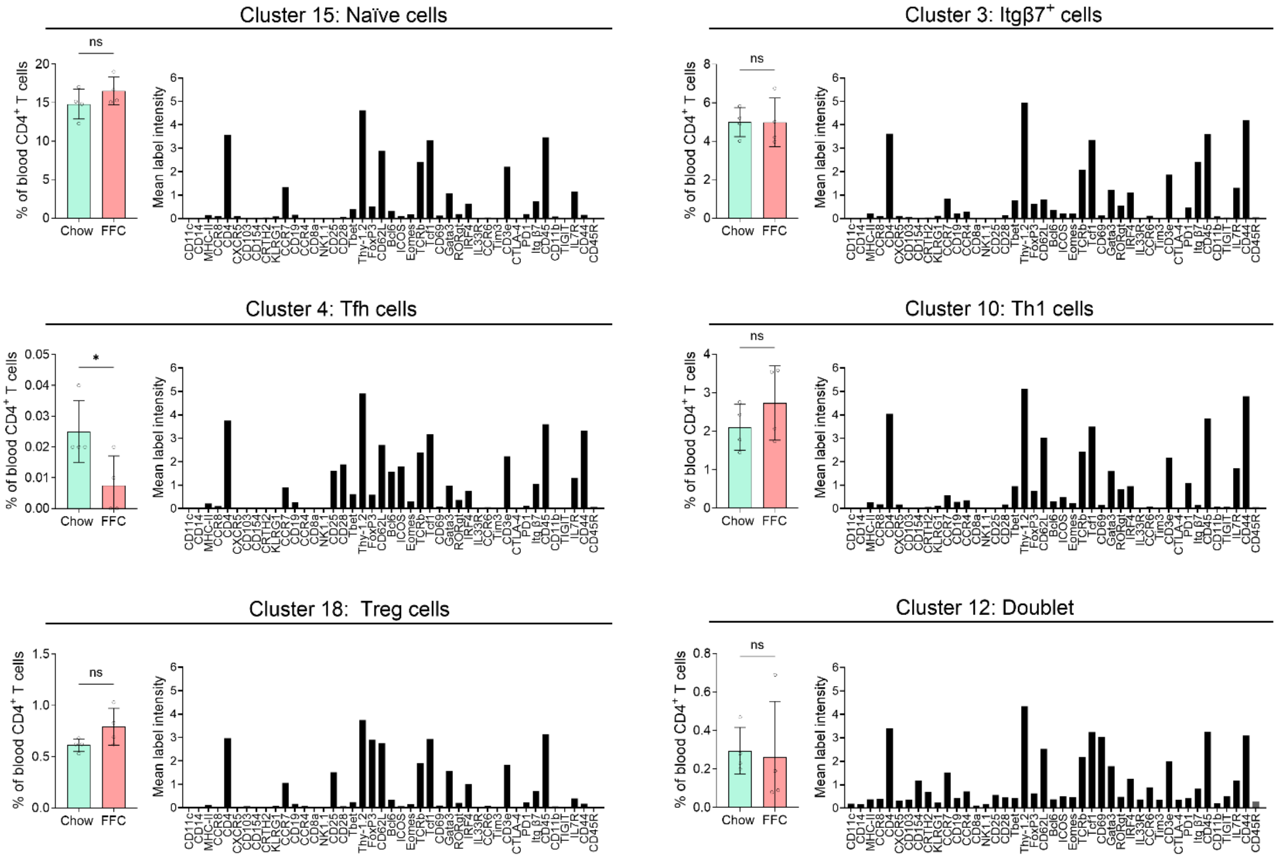


**Fig. S3. Peripheral blood and liver share similar changes in the CD4$^+$ T cell landscape during MASH. (A)** Mean label intensity for Th1, cytotoxic, Treg and Th17 cells showed in Figure 3. **(B)** Relative abundance and mean label intensity for remaining T cells clusters. Abbreviations: FFC, high fat, fructose, and cholesterol; PBMC, peripheral blood mononuclear cell; *p < 0.05, **p < 0.01, ***p < 0.001, ns, non-significant.

### Figure S4

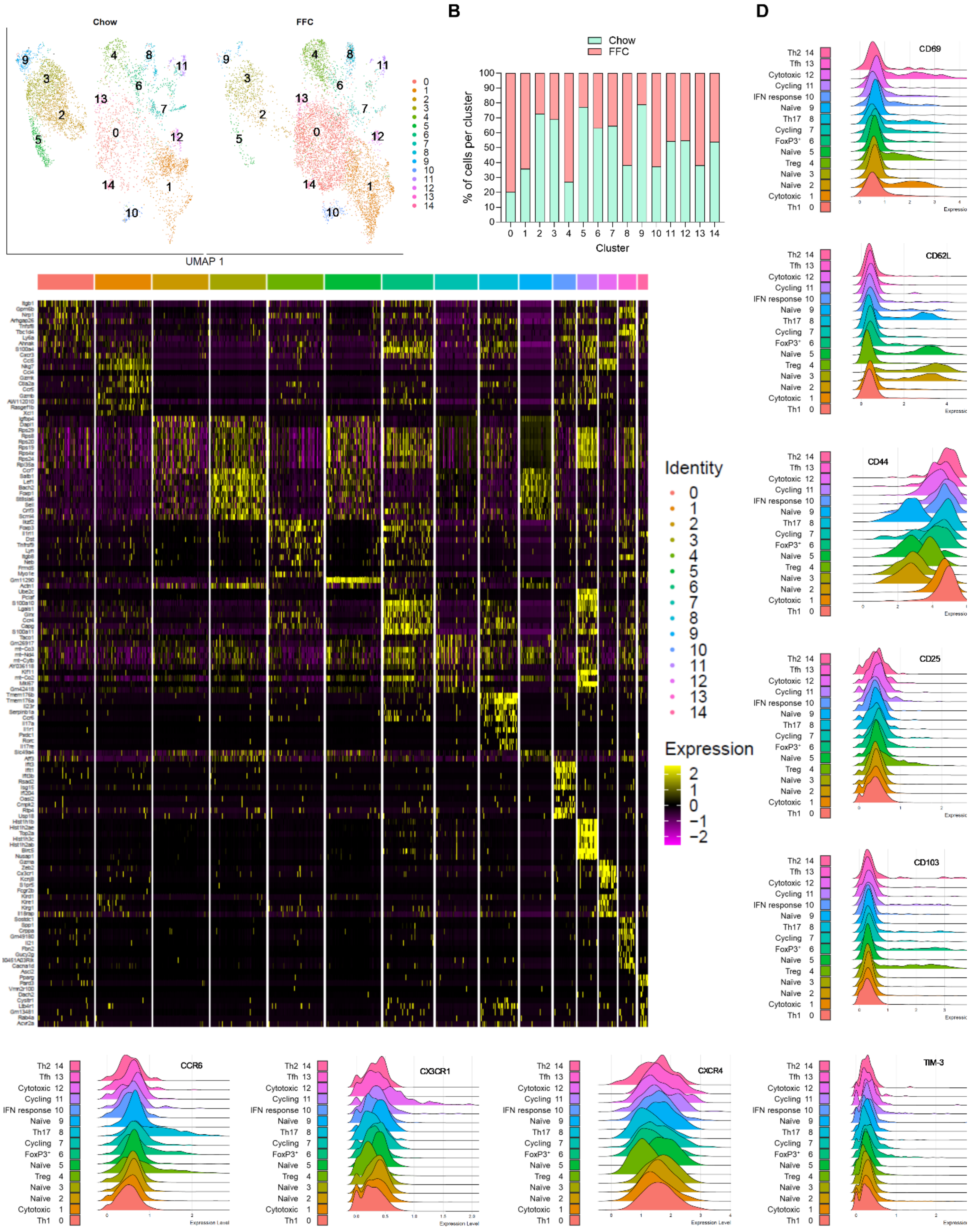

Chow
FFC
UMAP 1
B
Chow
FFC
% of cells per cluster
Cluster
D
Identity
Expression
CD69
CD62L
CD44
CD25
CD103
CCR6
CX3CR1
CXCR4
TIM-3
Th2 14
Tfh 13
Cytotoxic 12
Cycling 11
IFN response 10
Naïve 9
Th17 8
Cycling 7
FoxP3⁺ 6
Naïve 5
Treg 4
Naïve 3
Naïve 2
Cytotoxic 1
Th1 0
Expression Level

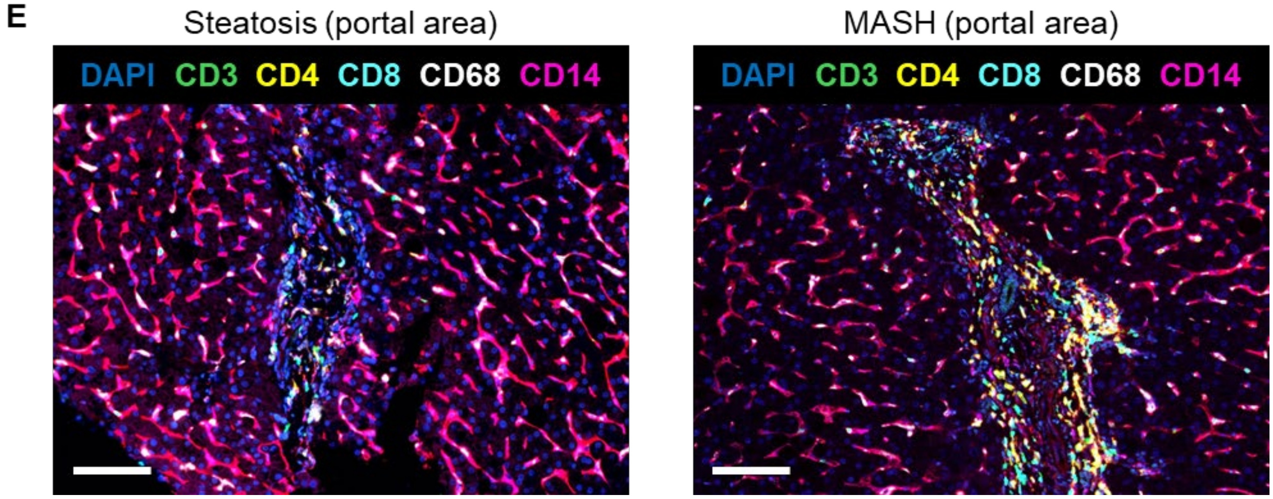


**Fig. S4. Single-cell atlas of intrahepatic CD4+ T cells in healthy and MASH livers. (A)** UMAP projection of hepatic $CD4^+$ T cells from chow and FFC-fed mice, showing 15 distinct clusters. **(B)** Percentage of cells within each cluster from chow versus FFC $CD4^+$ T cells. **(C)** Heatmap of top 10 differentially expressed genes across $CD4^+$ T-cell clusters. **(D)** Ridgeline plots showing relative expression of selected cell surface protein markers in $CD4^+$ T-cell clusters. **(E)** Representative images of human liver tissue from PhenoCycler. Abbreviations: UMAP, Uniform Manifold Approximation and Projection.

#### Figure S5

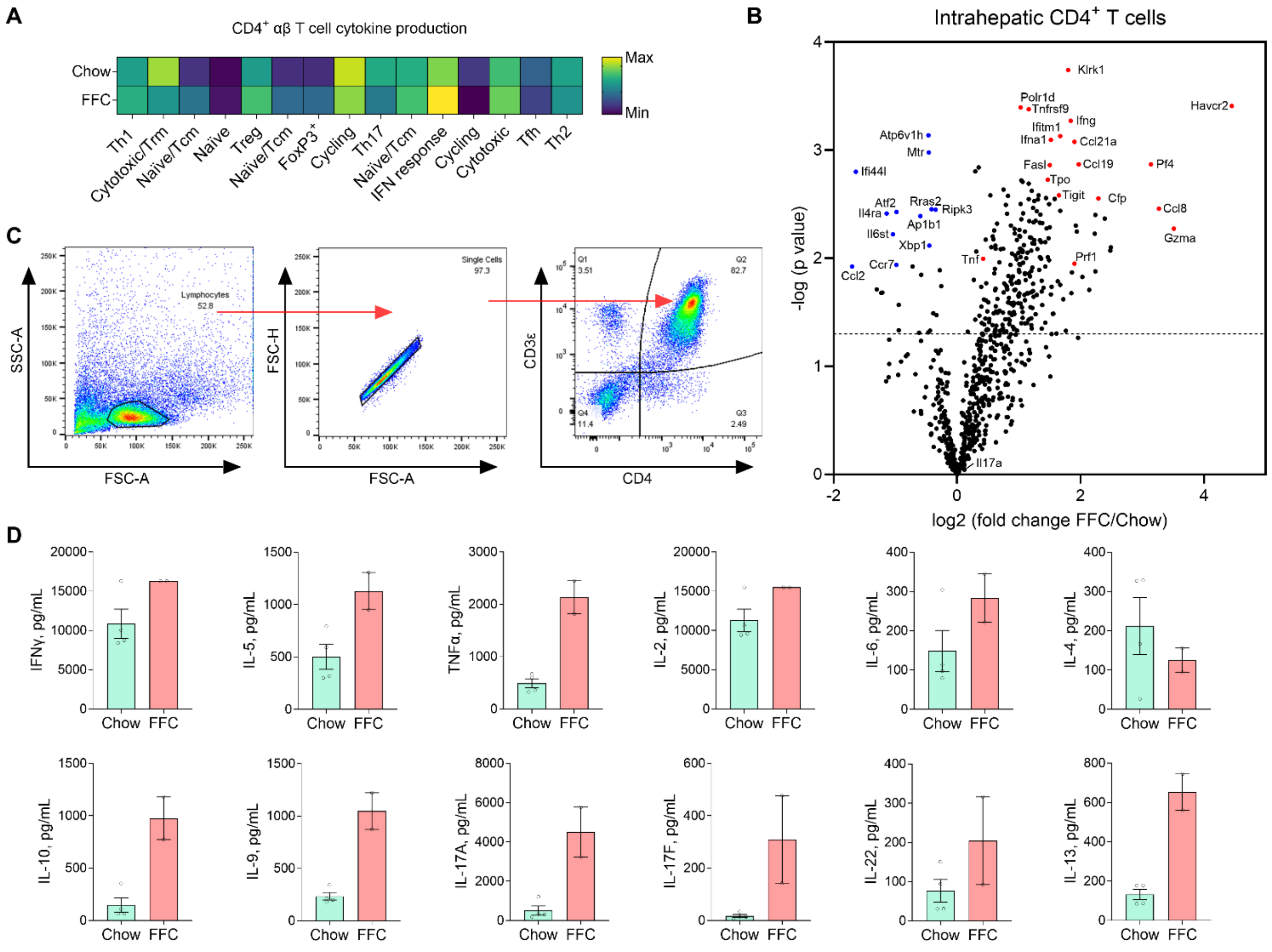


**Fig. S5. Liver CD4⁺ T cells in MASH display increased production of proinflammatory cytokines. (A)** Gene ontology biological process: CD4⁺ αβ T cell cytokine production pathway for CD4⁺ T cell clusters (chow vs FFC) obtained by CITEseq. Maximum and minimum values of enrichment score are shown in yellow and dark blue, respectively. **(B)** Total RNA from freshly isolated CD4⁺ T cells from chow and FFC livers was analyzed using a NanoString nCounter mouse autoimmunity panel with 770 mouse genes. Select upregulated genes are shown in red, and downregulated genes are shown in blue. **(C)** Gating strategy for assessment of intracellular cytokines in cultured hepatic CD4⁺ T cells. **(D)** CD4⁺ T cells from chow and FFC livers were re-stimulated with PMA and ionomycin. Cytokines secreted into cell culture supernatant were measured by LEGENDplex Mouse Th Cytokine Panel. N of samples indicated in the graphs. Abbreviations: SSC-A, side scatter area; FSC-A, forward scatter area; FSC-H, forward scatter height.

**Figure S6**

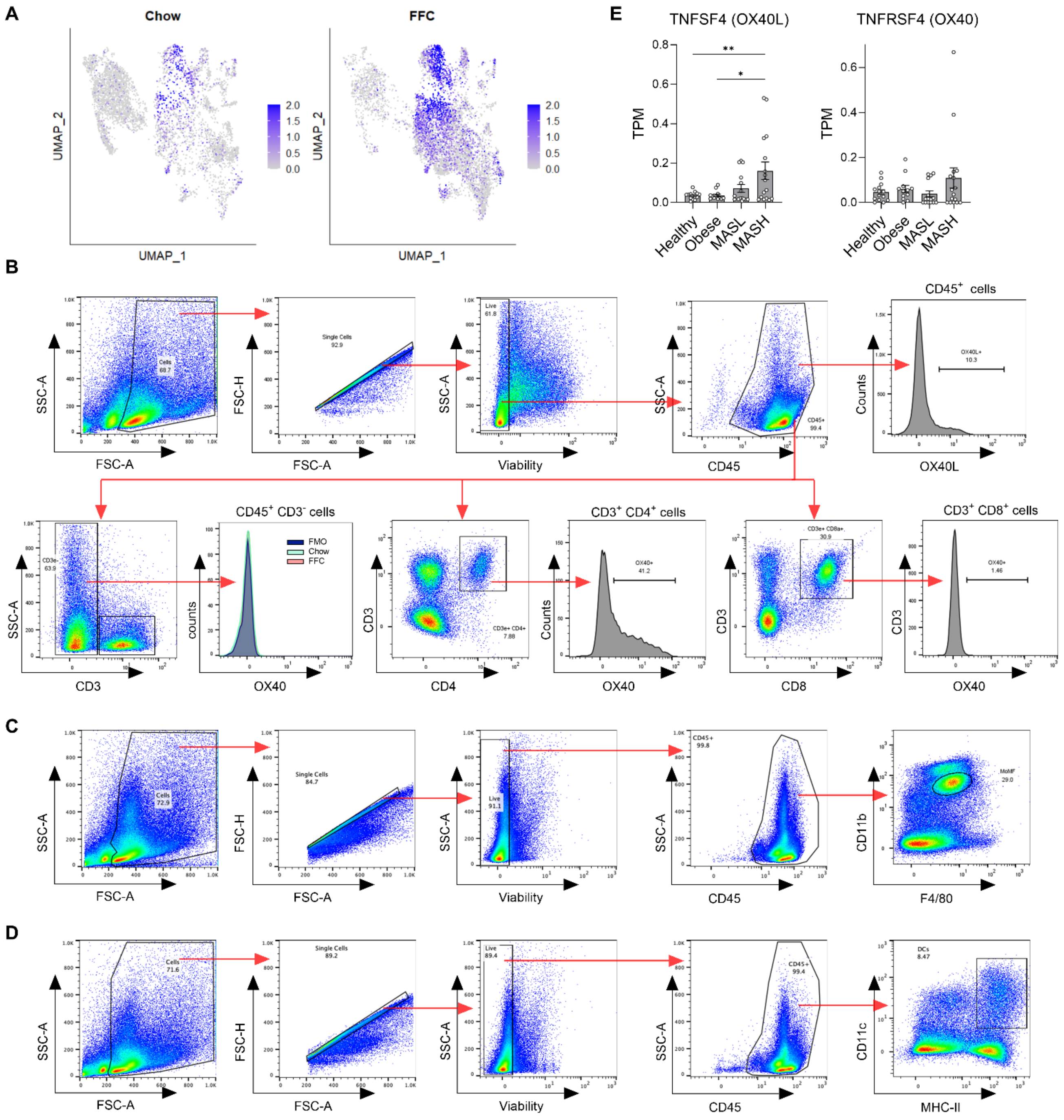


**Fig. S6: Tnfrsf4/OX40 is upregulated in CD4⁺ T cells in MASH. (A)** *Tnfrsf4* expression in murine hepatic CD4⁺ T cells by scRNAseq. **(B)** Gating strategy for assessing OX40 in hepatic CD4⁺ and CD8⁺ T cells and non-T cells by flow cytometry. **(C)** Gating strategy for OX40L expression in liver macrophages by flow cytometry. **(D)** Gating strategy for OX40L expression in liver dendritic cells by flow cytometry. **(E)** Gene expression of OX40L and OX40 in human liver tissues using a publicly available RNAseq dataset (GEO GSE126848). Representative gating is shown. Abbreviations: SSC-A, side scatter area; FSC-H, forward scatter height; TPM, Transcripts per kilobase million

#### Figure S7

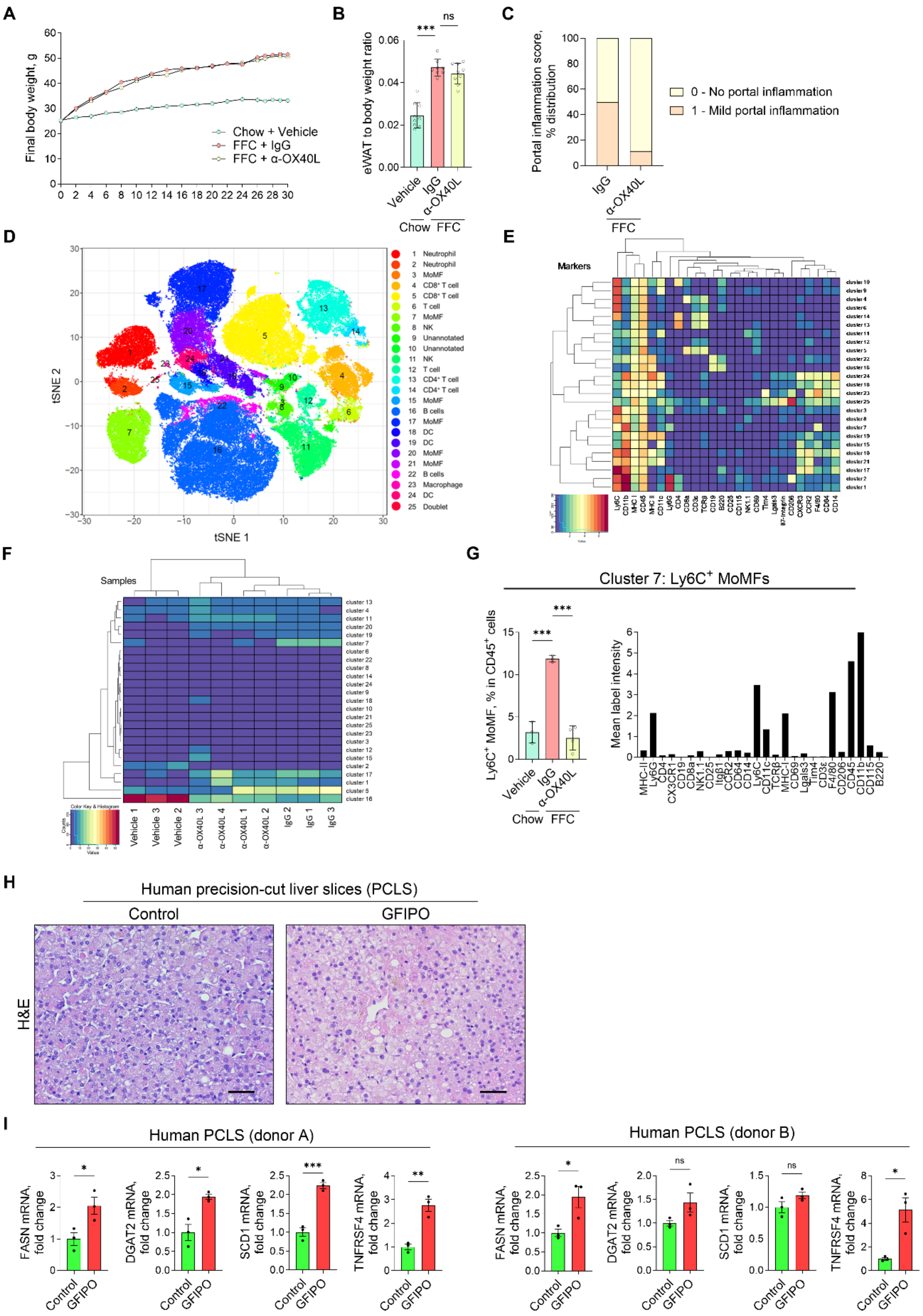
A
Final body weight, g
Chow + Vehicle
FFC + IgG
FFC + α-OX40L
B
eWAT to body weight ratio
Vehicle
IgG
α-OX40L
Chow
FFC
C
Portal inflammation score, % distribution
0 - No portal inflammation
1 - Mild portal inflammation
IgG
α-OX40L
FFC
D
tSNE 1
tSNE 2
E
Markers
F
Samples
G
Cluster 7: Ly6C⁺ MoMFs
Ly6C⁺ MoMF, % in CD45⁺ cells
Mean label intensity
H
Human precision-cut liver slices (PCLS)
Control
GFIPO
H&E
I
Human PCLS (donor A)
Human PCLS (donor B)
FASN mRNA, fold change
DGAT2 mRNA, fold change
SCD1 mRNA, fold change
TNFRSF4 mRNA, fold change
Control
GFIPO

**Fig. S7. Blocking the OX40L-OX40 axis reverses MASH and hepatic fibrosis in mice. (A)** Body weight time course during the 30-week study. **(B)** eWAT weight normalized to body weight at the time of sacrifice. **(C)** Portal inflammation score determined by a blinded pathologist. **(D)** Twenty-five unique clusters of cells were identified using an R-phenograph clustering algorithm, visualized on a tSNE plot, and annotated. **(E)** A heatmap of the relative intensity of protein markers across identified clusters with unsupervised hierarchical clustering. **(F)** Unsupervised hierarchical clustering of samples based on the relative abundance of individual clusters. **(G)** Relative abundance and mean label intensity of cluster 7, Ly6C$^{+}$ MoMF. **(H)** H&E staining of human PCLS incubated in control media or GFIPO media (containing glucose, fructose, insulin, palmitate, and oleate) for 96 hours. Representative images are shown; the scale bar is 50 µm. **(I)** Relative gene expression by qPCR in human PCLS incubated in control media or GFIPO media for 96 hours. Data points represent 3 slices per donor per condition. For panels B-G, N of samples is indicated in the graphs. ***p < 0.001. ns, non-significant. Abbreviations: FFC, high fat, fructose, and cholesterol; IgG, immunoglobulin G; eWAT, epidydimal white adipose tissue; tSNE, t-distributed stochastic neighbor embedding; MoMF, monocyte-derived macrophage; PCLS, precision-cut liver slices; GFIPO, glucose, fructose, insulin, palmitate, and oleate.

**Table S1. Antibody panel used for mass cytometric analysis of murine hepatic CD4⁺ T cells.**

| Label | Target | Clone | Manufacturer | Catalog # |
|---|---|---|---|---|
| 089Y | CD45 | 30-F11 | Fluidigm | 3089005B |
| 106Cd | CD14 | Sa14-2 | BioLegend | 123321 |
| 110Cd | I-A/I-E (MHC-II) | M5/114.15.2 | BioLegend | 107637 |
| 111Cd | CD198 (CCR8) | SA214G2 | BioLegend | 150302 |
| 112Cd | CD4 | RM4-5 | BioLegend | 100561 |
| 113Cd | CD185 (CXCR5) | L138D7 | BioLegend | 145502 |
| 114Cd | CD103 | 2E7 | BioLegend | 121402 |
| 116Cd | IL-21R | 4A9 | BioLegend | 131902 |
| 141Pr | ST2 (IL-33R) | DJ8 | MD Bioproducts | 101001 |
| 142Nd | EOMES | 1219A | R&D Systems | MAB8889 |
| 143Nd | TCRb | H57-597 | Fluidigm | 3143010B |
| 144Nd | TCF7/TCF1 | 812145 | R&D Systems | MAB8224 |
| 145Nd | CD69 | H1.2F3 | Fluidigm | 3145005B |
| 146Nd | GATA-3 | TWAJ | Thermo-Fisher | 14-9966-82 |
| 147Sm | CD196 (CCR6) | 29-2L17 | BioLegend | 129802 |
| 148Nd | ROR gamma (t) | B2D | Thermo-Fisher | 14-6981-82 |
| 149Sm | CD366 (Tim-3) | RMT3-23 | BioLegend | 119702 |
| 150Nd | IRF4 | IRF4.3E4 | BioLegend | 646402 |
| 151Eu | CD25 (IL-2R) | 3C7 | Fluidigm | 3151007B |
| 152Sm | CD3e | 145-2C11 | Fluidigm | 3152004B |
| 153Eu | CD28 | 37.51 | BioLegend | 102119 |
| 154Sm | CD152 (CTLA-4) | UC10-4B9 | Fluidigm | 3154008B |
| 155Gd | T-bet | 4B10 | BioLegend | 644825 |
| 156Gd | CD90.2/Thy-1.2 | 30-H12 | Fluidigm | 3156006B |
| 158Gd | FoxP3 | FJK-16s | Fluidigm | 3158003A |
| 159Tb | CD279 (PD-1) | RMP1-30 | Fluidigm | 3159006B |
| 160Gd | CD62L | MEL-14 | Fluidigm | 3160008B |
| 161Dy | CD154 (CD40L) | MR1 | Thermo Fisher | 16-1541-85 |
| 162Dy | CD294 (CRTH2) | No3m1scz | Invitrogen | CUST05491 |
| 163Dy | KLRG1 | 2F1 | BD Biosciences | 562190 |
| 164Dy | CD197 (CCR7) | 4B12 | Fluidigm | 3164013A |
| 165Ho | Bcl-6 | K112-91 | BD Biosciences | 561520 |
| 166Er | CD19 | 6D5 | Fluidigm | 3166015B |
| 167Er | CD194 (CCR4) | 2G12 | BioLegend | 131202 |
| 168Er | CD8a | 53-6.7 | Fluidigm | 3168003B |
| 169Tm | Integrin β7 | FIB504 | BioLegend | 321202 |
| 170Er | CD161 (NK1.1) | PK136 | Fluidigm | 3170002B |
| 171Yb | CD11b | M1/70 | BioLegend | 101249 |
| 173Yb | TIGIT (Vstm3) | 1G9 | BioLegend | 142101 |
| 174Yb | CD127 (IL-7Ra) | A7R34 | Fluidigm | 3174013B |
| 175Lu | CD278 (ICOS) | C398.4A | Fluidigm | 3148019B |
| 176Yb | CD44 | IM7 | BioLegend | 103051 |
| 198Pt | CD6 | OX-129 | BioLegend | 146404 |
| 209Bi | CD11c | N418 | Fluidigm | 3209005B |

**Table S2. Antibody panel used for mass cytometric analysis of murine PBMCs.**

| Label | Target | Clone | Manufacturer | Catalog # |
|---|---|---|---|---|
| 089Y | CD45 | 30-F11 | Fluidigm | 3089005B |
| 106Cd | CD14 | Sa14-2 | BioLegend | 123321 |
| 110Cd | I-A/I-E (MHC-II) | M5/114.15.2 | BioLegend | 107637 |
| 111Cd | CD198 (CCR8) | SA214G2 | BioLegend | 150302 |
| 112Cd | CD4 | RM4-5 | BioLegend | 100561 |
| 113Cd | CD185 (CXCR5) | L138D7 | BioLegend | 145502 |
| 114Cd | CD103 | 2E7 | BioLegend | 121402 |
| 116Cd | CD45R (B220) | RA3-6B2 | BioLegend | 103249 |
| 141Pr | ST2 (IL-33R) | DJ8 | MD Bioproducts | 101001 |
| 142Nd | EOMES | 1219A | R&D Systems | MAB8889 |
| 143Nd | TCRb | H57-597 | Fluidigm | 3143010B |
| 144Nd | TCF7/TCF1 | 812145 | R&D Systems | MAB8224 |
| 145Nd | CD69 | H1.2F3 | Fluidigm | 3145005B |
| 146Nd | GATA-3 | TWAJ | Thermo-Fisher | 14-9966-82 |
| 147Sm | CD196 (CCR6) | 29-2L17 | BioLegend | 129802 |
| 148Nd | ROR gamma (t) | B2D | Thermo-Fisher | 14-6981-82 |
| 149Sm | CD366 (Tim-3) | RMT3-23 | BioLegend | 119702 |
| 150Nd | IRF4 | IRF4.3E4 | BioLegend | 646402 |
| 151Eu | CD25 (IL-2R) | 3C7 | Fluidigm | 3151007B |
| 152Sm | CD3e | 145-2C11 | Fluidigm | 3152004B |
| 153Eu | CD28 | 37.51 | BioLegend | 102119 |
| 154Sm | CD152 (CTLA-4) | UC10-4B9 | Fluidigm | 3154008B |
| 155Gd | T-bet | 4B10 | BioLegend | 644825 |
| 156Gd | CD90.2/Thy-1.2 | 30-H12 | Fluidigm | 3156006B |
| 158Gd | FoxP3 | FJK-16s | Fluidigm | 3158003A |
| 159Tb | CD279 (PD-1) | RMP1-30 | Fluidigm | 3159006B |
| 160Gd | CD62L | MEL-14 | Fluidigm | 3160008B |
| 161Dy | CD154 (CD40L) | MR1 | Thermo Fisher | 16-1541-85 |
| 162Dy | CD294 (CRTH2) | No3m1scz | Invitrogen | CUST05491 |
| 163Dy | KLRG1 | 2F1 | BD Biosciences | 562190 |
| 164Dy | CD197 (CCR7) | 4B12 | Fluidigm | 3164013A |
| 165Ho | Bcl-6 | K112-91 | BD Biosciences | 561520 |
| 166Er | CD19 | 6D5 | Fluidigm | 3166015B |
| 167Er | CD194 (CCR4) | 2G12 | BioLegend | 131202 |
| 168Er | CD8a | 53-6.7 | Fluidigm | 3168003B |
| 169Tm | Integrin β7 | FIB504 | BioLegend | 321202 |
| 170Er | CD161 (NK1.1) | PK136 | Fluidigm | 3170002B |
| 171Yb | CD11b | M1/70 | BioLegend | 101249 |
| 173Yb | TIGIT (Vstm3) | 1G9 | BioLegend | 142101 |
| 174Yb | CD127 (IL-7Ra) | A7R34 | Fluidigm | 3174013B |
| 175Lu | CD278 (ICOS) | C398.4A | Fluidigm | 3148019B |
| 176Yb | CD44 | IM7 | BioLegend | 103051 |
| 209Bi | CD11c | N418 | Fluidigm | 3209005B |

**Table S3. Antibody panel used for mass cytometric analysis of murine liver leukocytes.**

| Label | Target | Clone | Manufacturer | Catalog # |
|---|---|---|---|---|
| 089Y | CD45 | 30-F11 | Fluidigm | 3089005B |
| 141Pr | Lgals3 | 202213 | R&D Systems | MAB1197 |
| 142Nd | CD11c | N418 | Fluidigm | 3142003B |
| 143Nd | TCRb | H57-597 | Fluidigm | 3143010B |
| 144Nd | MHC Class I | 28-14-8 | Fluidigm | 3144016B |
| 145Nd | CD69 | H1.2F3 | Fluidigm | 3145005B |
| 149Sm | Tim-4 | RMT4-54 | BioLegend | 130002 |
| 151Eu | CD25 (IL-2R) | 3C7 | Fluidigm | 3151007B |
| 152Sm | CD3e | 145-2C11 | Fluidigm | 3152004B |
| 153Eu | CD29 | 9EG7 | BD Biosciences | 550531 |
| 156Gd | CCR2 | 475301 | R&D Systems | MAB55381 |
| 159Tb | F4/80 | BM8 | Fluidigm | 3159009B |
| 160Gd | CD64 | 290322 | R&D Systems | MAB20741 |
| 161Dy | Ly-6G | 1A8 | BioLegend | 127637 |
| 163Dy | CD4 | RM4-5 | BioLegend | 100561 |
| 164Dy | CX3CR1 | SA011F11 | Fluidigm | 3164023B |
| 165Ho | CD14 | Sa14-2 | BioLegend | 123321 |
| 166Er | CD19 | 6D5 | Fluidigm | 3166015B |
| 168Er | CD8a | 53-6.7 | Fluidigm | 3168003B |
| 169Tm | CD206 (MMR) | C068C2 | Fluidigm | 3169021B |
| 170Er | CD161 (NK1.1) | PK136 | Fluidigm | 3170002B |
| 171Yb | CD11b | M1/70 | BioLegend | 101249 |
| 174Yb | CD115 (CSF-1R) | AFS98 | BioLegend | 135521 |
| 175Lu | Ly-6C | HK1.4 | BioLegend | 128039 |
| 176Yb | CD45R (B220) | RA3-6B2 | Fluidigm | 3176002B |
| 209Bi | I-A/I-E (MHC-II) | M5/114.15.2 | Fluidigm | 3209006B |

**Table S4. Antibodies used for flow cytometry and cell sorting**

| Target | Fluorochrome | Clone | Manufacturer | Catalog # |
|---|---|---|---|---|
| CD3e | AF488 | 500A2 | BioLegend | 152322 |
| CD3e | PE-Cy7 | 500A2 | BioLegend | 152313 |
| CD4 | PE/Cy7 | GK1.5 | BioLegend | 100421 |
| CD4 | AF647 | RM4-5 | BioLegend | 100530 |
| CD4 | PerCP-Vio 700 | REA1211 | Miltenyi Biotec | 130-123-213 |
| IFNg | BV421 | XMG1.2 | BioLegend | 505830 |
| TNFa | PE | MP6-XT22 | BioLegend | 506305 |
| IL-17A | PE/Cy7 | TC11-18H10.1 | BioLegend | 100421 |
| CD8a | APC-Cy7 | 53-6.7 | BioLegend | 100714 |
| CD134 (OX40) | BV-421 | OX-86 | BioLegend | 119411 |
| PBS-57 loaded CD1d tetramer | PE | N/A | NIH Tetramer Core Facility | N/A |
| CD1d tetramer (unloaded) | PE | N/A | NIH Tetramer Core Facility | N/A |
| CD45 | VioGreen | REA737 | Miltenyi Biotec | 130-110-803 |
| CD11b | AF488 | M1/70 | BioLegend | 101219 |
| F4/80 | PE-Cy7 | BM8 | BioLegend | 123113 |
| OX40L | APC | REA960 | Miltenyi Biotec | 130-116-073 |
| CD11C | FITC | N418 | BioLegend | 117305 |
| MHC II | PE | REA813 | Miltenyi Biotec | 130-112-387 |
| CD11b | PerCP-Vio700 | REA592 | Miltenyi | 130-113-809 |
| Ly6G | PE | REA526 | Miltenyi | 130-123-780 |
| F4/80 | PE/Cy7 | BM 8 | BioLegend | 123113 |

**Table S5. CITE-seq antibody panel for murine hepatic CD4$^+$ T cells**

| Target | Clone | Barcode Sequence | Manufacturer | Catalog # |
|---|---|---|---|---|
| CD366 (Tim-3) | RMT3-23 | ATTGGCACTCAGATG | BioLegend | 119741 |
| CD44 | IM7 | TGGCTTCAGGTCCTA | BioLegend | 103071 |
| CD25 | PC61 | ACCATGAGACACAGT | BioLegend | 102067 |
| CD62L | MEL-14 | TGGGCCTAAGTCATC | BioLegend | 104465 |
| CD69 | H1.2F3 | TTGTATTCCGCCATT | BioLegend | 104555 |
| CD127 (IL-7Ra) | A7R34 | GTGTGAGGCACTCTT | BioLegend | 135055 |
| CD103 | 2E7 | TTCATTAGCCCGCTG | BioLegend | 121445 |
| Integrin β7 | FIB504 | TCCTTGGATGTACCG | BioLegend | 321231 |
| CD196 (CCR6) | 29-2L17 | CTCTCTGCATTCCTC | BioLegend | 129827 |
| CD183 (CXCR3) | CXCR3-173 | GTTCACGCCGTGTTA | BioLegend | 126549 |
| CD197 (CCR7) | 4B12 | TTATTAACAGCCCAC | BioLegend | 120133 |
| CD184 (CXCR4) | L276F12 | GTCGTGGTGTTGTTC | BioLegend | 146521 |
| CD304 (NRP1) | 3E12 | CCAGCTCATTCAACG | BioLegend | 145219 |
| CX3CR1 | SA011F11 | CACTCTCAGTCCTAT | BioLegend | 149045 |
| CD194 (CCR4) | 2G12 | TTCATGTGTTTGTGC | BioLegend | 131223 |
| CD185 (CXCR5) | L138D7 | ACGTAGTCACCTAGT | BioLegend | 145539 |
| CD198 (CCR8) | SA214G2 | ATCTCCGTTGTGCGA | BioLegend | 150331 |

**Table S6. Antibody panel for multiplexed spatial phenotyping using PhenoCycler (Akoya)**

| Target | Clone | Manufacturer | Catalog # |
|---|---|---|---|
| CD11c | D3V1E | Cell Signaling Technology | 45581 |
| HLA-E | MEM-E/02 | Abcam | ab2216 |
| CD163 | DGU1J | Cell Signaling Technology | 25121SF |
| CD57 | HNK-1 | BioLegend | 359602 |
| GATA3 | D13C9 | Cell Signaling Technology | 10630SF |
| Podoplanin | NC-08 | BioLegend | 337002 |
| CD31 | EP3095 | Abcam | ab226157 |
| CD4 | EPR6855 | Abcam | ab181724 |
| HLA-A | EP1395Y | Abcam | ab216653 |
| CD20 | L26 | Thermo Fisher | MA513141 |
| CD40 | D8W3N | Cell Signaling Technology | 77841SF |
| SMA | 1A4 | Abcam | ab240654 |
| E-cadherin | 4A2C7 | Thermo Fisher | 33-4000 |
| CD68 | KP1 | Abcam | ab233172 |
| CD66 | ASL-32 | BioLegend | 342302 |
| CD45RO | UCH-L1 | Abcam | ab23 |
| Pan-cytokeratin | AE-1/AE-3 | Abcam | ab80826 |
| CD38 | E7Z8C | Cell Signaling Technology | 43382SF |
| CD45 | D9M8I | Cell Signaling Technology | 47937SF |
| Vimentin | O91D3 | BioLegend | 677802 |
| Myeloperoxidase | E1E7I | Cell Signaling Technology | 88757SF |
| CD34 | QBEnd/10 | Abcam | ab213054 |
| CD8a | C8/144B | BioLegend | 372902 |
| IDO | V1NC3IDO | Thermo Fisher | 14-9750-82 |
| CD39 | EPR20627 | Abcam | ab236038 |
| FoxP3 | 236A/E7 | Abcam | ab96048 |
| CD21 | EP3093 | Abcam | ab271855 |
| HLA-DR | EPR3692 | Abcam | ab209968 |
| Galectin 3 | M3/38 | BioLegend | 125401 |
| CD14 | EPR3653 | Abcam | ab226121 |
| Granzyme B | D6E9W | Cell Signaling Technology | 79903SF |
| Collagen IV | EPR20966 | Abcam | ab226485 |
| PD-L1 | 73-10 | Abcam | ab226766 |
| CD3e | EP449E | Abcam | ab271850 |
| PD1 | D4W2J | Cell Signaling Technology | 63815SF |
| Ki-67 | B56 | Abcam | ab279657 |
| TOX | E6I3Q | Cell Signaling Technology | 62886SF |
| T-bet/TBX21 | D6N8B | Cell Signaling Technology | 27112SF |
| ICOS | D1K2T | Cell Signaling Technology | 39740SF |
| LAG-3 | EPR20261 | Abcam | ab227579 |
| Na/K ATPase | EP1845Y | Abcam | ab167390 |
| CD79a | D1X5C | Cell Signaling Technology | 84162SF |
| EpCAM | D9S3P | Cell Signaling Technology | 55725SF |
| Eomes | D8D1R | Cell Signaling Technology | 71103 |
| RORgt | NA | Thermo-Fisher | PA5-23148 |

**Table S6. Primer sequences for qPCR**

| Target | Forward primer (5'-3') | Reverse primer (5'-3') |
|---|---|---|
| 18S | CGCTTCCTTACCTGGTTGAT | GAGCGACCAAAGGAACCATA |
| | Murine | |
| Spp1 | CTCCATCGTCATCATCATCG | TGCACCCAGATCCTATAGCC |
| Trem2 | CTGGAACCGTCACCATCACTC | CGAAACTCGATGACTCCTCGG |
| Cd9 | ATGCCGGTCAAAGGAGGTAG | GCCATAGTCCAATAGCAAGCA |
| Fabp5 | TGAAAGAGCTAGGAGTAGGACTG | CTCTCGGTTTTGACCGTGATG |
| Gpnmb | TGCCAAGCGATTTCGTGATGT | GCCACGTAATTGGTTGTGCTC |
| | Human | |
| IFNG | TCGGTAACTGACTTGAATGTCCA | TCGCTTCCCTGTTTTAGCTGC |
| CXCL10 | GTGGCATTCAAGGAGTACCTC | GCCTTCGATTCTGGATTCAG |
